# Impact of glacier retreat on biodiversity and plant–insect interaction dynamics

**DOI:** 10.1101/2023.04.22.537665

**Authors:** Bao Ngan Tu, Nora Khelidj, Gianalberto Losapio

## Abstract

With the retreat and extinction of glaciers worldwide, new areas are exposed for colonization by diverse plants and associated insects. Yet, glacier retreat is also followed by the loss of plants and insects from local communities, causing changes in species diversity, species composition and plant–insect interactions. However, the impact of glacier retreat and extinction on pollination networks remains poorly understood. An integrative understanding of pollination network dynamics following glacier retreat is therefore of major importance to biodiversity maintenance and ecosystem functioning and services. Here, we addressed how glacier retreat affects directly and indirectly through biodiversity the frequency, complexity, and diversity of plant–insect interactions. After reconstructing the geochronology of glaciers (Mont Miné glacier, Swiss Alps), we surveyed plant–insect interactions and analyzed network dynamics. We observed sharp changes in the diversity of both plant and insect communities. We found an increase in the frequency of their interactions following glacier retreat, but an ultimate decrease with glacier extinction. Yet, after controlling for the effects of flower diversity, interaction frequency showed a regular, ‘universal’ pattern. Accordingly, the complexity of pollination networks and interaction diversity tended to change at constant rates with glacier retreat. Our results indicate that, in the long-term, glacier retreat decreases biodiversity and influence the stability of ecological networks. The good news is that increasing flower diversity would counteract these impacts by increasing interaction diversity and complexity. Supporting plant and flower diversity may therefore be a key strategy for halting the erosion of ecological networks while increasing ecosystem functioning.

## INTRODUCTION

### 1. Glacier retreat and colonization

The retreat and extinction of glaciers worldwide is a visible proof of global warming’s effects (Marzeion et al., 2014; Zemp et al., 2015; Roe et al., 2017). We are now facing the extinction of glaciers at unprecedent rate since the early 21st century (Zemp et al., 2015, 2019) from tropical and Mediterranean mountains (Apennine, Sierra Nevada) to the European Alps, Andes, Caucasus and Himalayas (Huss et al., 2017; Sommer et al., 2020). In the European Alps, the Little Ice Age (LIA, from the mid-1200s) was the main period of glaciers re-advance with the latest maxima in the early and mid-1800s (Nicolussi et al., 2022). Since the LIA maxima (approximately 1850), glaciers in European Alps are retreating continuously with a speed now 200-300% faster than it was 20 years ago and already have lost 25-30% of surfaces (Sommer et al., 2020; Marta et al., 2021).

Glacier retreat affects landscape composition, soil properties, water resources and micro- and macroclimate (Fell et al., 2017; Brighenti et al., 2019; Cauvy-Fraunié and Dangles, 2019; Ficetola et al, 2021). The glacier melting exposes the new arid ice-free areas such as forelands (areas in front of retreating glaciers) (Matthews 1992). Over time, forelands are continuously revealed and colonized by diverse living organisms, then complex and functional ecosystems are formed. Plants are known as the primary first pioneers (Whittaker 1993; Chapin et al., 1994; Tampucci et al., 2015; Anthelme et al., 2021). For instance, in the early stage of high tropical alpine glacier retreat, lichen and bryophytes were dominant and played a key role as pioneers while vascular plants covered were lower for around 60 years (Llambí et al., 2021). Following plants, others living organisms such as invertebrates and vertebrates occurred to take advantages from new habitat (Tampucci et al., 2015; Junker et al., 2020). This colonization process is resulting in younger communities present in front of the current glacier while the older communities become more distant (Ficetola et al., 2021; Hanusch et al., 2022;). The fact that glacier retreat creates visible glacier path (which is moraine) in between two areas. Using evidence provided by geochronology reports, one can reconstruct the glacier retreat’s phases in a chronosequence, then calculate the age (the time passed since the glacier retreat) for the terrains and their relevant communities (Losapio et al., 2021a; Ficetola et al., 2021). For these reasons, we can assume that time since deglaciation plays an important role in the evolution of ecosystems (Whittaker, 1993; Ficetola et al., 2021).

Noticeably, this process is followed by distribution shifts and creates new interactions among species (Cauvy-Fraunié and Dangles, 2019; Ficetola et al, 2021). The geo-ecological model (Whittaker, 1993) assumes that in the initial stages of colonization, environmental conditions are more important than organisms in controlling community development. While, in later stages, the modification of habitat influenced by interactions between organisms and effects of neighbor plants in altering biodiversity becomes more important (Erschbamer and Caccianiga, 2016; Losapio et al., 2021b). In addition, glacier extinction will also be followed by the loss of species and erosion of ecological networks in local communities (Dullinger et al., 2012; Losapio et al., 2015, 2021a). Especially, plants and insects play an important role in the planet ecosystem as mutually relative organisms, but they are highly sensitive to the change of environment (Erschbamer and Caccianiga, 2016; Ficetola et al, 2021; Losapio et al., 2015, 2021a).

### 2. Plant diversity

As the primary first pioneers, plant communities are strongly influenced by glacier retreat (Whittaker 1993; Chapin et al., 1994; Anthelme et al., 2021). Plants play an important role as the primary producers that created the base of the food web that maintains other organisms’ lives, and plant diversity is also a major key to healthy ecosystems (Inouye, 2020). Alpine environments are biodiversity hotsposts and habitats which are extremely sensitive to climate change (Cauvy-Fraunié and Dangles, 2019). Scientists give one prediction about the European mountain ecosystem: 36–55% of alpine species, 31–51% of subalpine species, and 19– 46% of mountain species will lose more than 80% of their suitable habitat by 2070–2100 (Inouye, 2020). The retreat of glaciers in the European Alps pushes their unique plant communities to face adaptation or extinction. Some studies show evidence that biodiversity increases with glacier retreat at a local level in the short term. For instance, Cauvy-Fraunié and Dangles determined an increase in the abundance and richness of generalist taxa but a decrease in specialist species. They concluded that these “winners” and “losers” presented the local increase of biodiversity as glacier retreat (Cauvy-Fraunié and Dangles, 2019). In addition, when studying the effects of glacier retreat on plant species, Losapio et al., found an initial increase in plant diversity under glacier retreat (Losapio et al., 2021a). However, following the global extinction of glaciers, total biodiversity will be influenced negatively (Stibal et al., 2020).

In fact, we observed the ultimate decrease in plant diversity after glacier extinction (Losapio et al., 2021a). In the study of glacier retreat’s impacts on plant primary succession (2-13 years after glacier retreat) through plant diversity, plant traits, and microenvironment, scientists found a decrease in both plant population level (lower plant height) and the community levels (lower species richness and abundance) (Anthelme et al., 2021). Additionally, the results when studying the emergence of multi-diversity in Ödenwinkel glacier (Austria) reported a clear increase in plant diversity (Shannon index) in the first 110 years after retreat, then decrease after succession proceeds (Junker et al., 2020). Also, not all species are equal under global warming, some species gain benefits and become “the winner” while others called “the loser” face struggle and even extinction (Losapio et al., 2016; Cauvy-Fraunié and Dangles, 2019).

When doing researches of which factors drive plant diversity in glacier foreland, complex combinations of negative and positive biotic impact structures of plant communities were provided (Whittaker, 1993; Chapin et al., 1994). Ficetola et al. reviewed three patterns of community dynamics: (1) additions, (2) replacements, and (3) tipping points. The addition model assumes the important role of differences in colonization or tolerance while the replacements model emphasizes the competition between groups. These two patterns can happen together and are distinguished by the tipping points where the limitation and competition become stronger (Ficetola et al. in 2021). Plant communities often follow the replacement model, presenting a major role of competition in favorable environments (later stages) (Chapin et al. 1994; Ficetola et al, 2021). This dynamic was supported by the results from Losapio et al., 2021 indicating the shifts within plant communities: in the long term, competitive species become more predominant while some of the first colonization species, such as *Artemisia genipi* were declined within only 100 years. Thus, the local extinction of those plants can also lead to the risk of others species and affect the functional ecosystem (Losapio et al., 2021a).

### 3. Insect diversity

Next to plants, the following colonizers in a successional process after glacier retreat are invertebrates (Tampucci et al., 2015; Junker et al., 2020). Here we are interested in invertebrates, especially pollinator insects which have a close, mutual relationship with plants in the Alpine glacier foreland. Without pollinators, we would face a large decrease in pollinated-insect plants (Losapio et al., 2016; Adedoja et al., 2018; Inouye, 2020; Hanusch et al., 2022).

Pollinators have been studied in the Alpine ecosystem since the 1800s. Scientists have noted the presence of bees, butterflies, moths, beetles or ants that are characteristic of plant pollination at high altitudes in the Alps (Inouye, 2020). They observed the shift in pollinator communities following altitude gradient. For instance, in areas where social bumblebees (Bombus sp.) are not found, flies (syrphid flies) and short-tongued solitary bees have taken the place and become the most important high-altitude pollinators or take more advances than bees on pollination at lower elevations (McCabe and Cobb, 2021). Besides, insects are found as pioneers without the presence of plants in surface of glaciers, glacial landforms and recent glaciers (Hågvar, 2012; Hågvar et al., 2020; Gobbi and Lencioni, 2021). Arthropods were found increasing in the begin after glacier retreat but slightly decreases on the oldest plots but later than plant communities about 40 years (Junker et al., 2020). In contrast, at high-elevation mountains, the loss of habitat (especially ice-related landforms) and global warming bring negative effects to the cold-adapted insect species community. The loss (ca. 9%) of terrestrial insect abundance per decade over space and time was reported (Gobbi and Lencioni, 2021).

With three patterns of community dynamics mentioned above (Ficetola et al., 2021), scientists found that in replacement model, some invertebrates communities were driven mainly by biotic relations (i.e., plant–insect interactions) (Kaufmann & Raffl 2002). Conversely, in addition pattern, the differences within insect community’s structure were more important (Gobbi et al. 2021; Ficetola et al., 2021). That evidence indicates that besides the impact of abiotic factors, the interaction between communities and also their traits can drive the insect community dynamics.

### 4. Plant–insect interaction network

Like the two components of the interaction, plant–insect interaction networks are very sensitive to the changes in environmental conditions as well as the distribution of interacting partners (Adedoja et al., 2018; Bonelli et al., 2022). Plant-insect interactions are generally complex and reflect the functioning and health of the ecosystem (Adedojia et al., 2018; Losapio et al., 2020; Badenes-Pérez, 2022; Bonelli et al., 2022). The relationship between plants and insects has different components. Insects may act as protection, dispersers, or pollinators for plants while plants may be a food resource or nest location for insects (Calatayud et al., 2018). Yet, both partners benefit from this relationship.

Although plants and insects are both among the primary pioneers in ecological succession, research on plant and insect communities along glacier retreats were studied independently and tended to focus on plants (Whittaker 1993; Chapin et al., 1994; Dullinger et al., 2012; Trunschke and Sto klin, 2017; Llambí et al., 2021; Bonelli et al., 2022; Brochmann et al., 2022). More recent studies on ecological succession after glacier retreat have supplemented advances in research of community dynamics by using distribution, functional, and network theories (Ficetola et al., 2021). Since then, recent researches have gradually taken interest in studying the role of biological interactions in shaping biodiversity, leading species distributions and supplying species responses under the influence of glacier retreat (Losapio et al., 2015, 2016, 2019, 2021b; Ershbamer and Caccianiga, 2016; Valle et al., 2022a, b). For instance, Losapio’s study reported an increase of plant–anthophilous insect interactions and the shifts of the network from pollinator to diverse food web along the glacier retreat (Losapio et al., 2015). When going further on studying feedback effects between plants and pollinators, they determined that the dynamics of alpine plant and insect communities may be driven by biotic interactions and feedback processes more than just the abiotic conditions (Losapio et al., 2016). Furthermore, plant community was a stronger predictor of network complexity and interaction diversity than the insect community (Robinson et al., 2018). Additionally, Bonelli et al. reported the effect of temperature and time (hour of the day), but not wind speed and the number of flowers per plant on the flower-visiting of arthropods (Bonelli et al., 2022). Yet, it is not clear how interaction diversity changes with glacier retreat.

Indeed, the lack of comprehensive network-level studies impairs our ability to predict the fate of biodiversity and the functioning of ecosystems on glacier foreland. An integrative understanding of how plant and insect communities and their network dynamics respond following glacier retreat is therefore of major importance to biodiversity maintenance and environmental health (Erschbamer and Caccianiga, 2016; Giron et al., 2018; Ficetola et al, 2021). We hypothesize that glacier retreat and extinction will decrease plant and insect diversity and their interaction diversity. Furthermore, we expect the strong effect of time (years after glacier retreat) and plant richness on driving community dynamics. In particular, we addressed the following questions: (i) How does glacier retreat affect the diversity of plant and insect communities? (ii) How do plant–insect interactions change in response to glacier retreat (iii) Does plants contribute to the change of insect diversity and plant–insect interaction networks under the glacier retreat?

## MATERIALS AND METHODS

### 1. Study site

The study was performed along the foreland of Mont Miné glacier in the Hérens valley (central Swiss Alps, 46°01′ N/7°33′ E). Mont Miné glacier covers an area of 9.67km^2^. The highest and lowest point of the glacier were 3804 and 2122 m a.s.l. in 2015 (Paul et al., 2020; Nicolussi et. al., 2022). After the last Little Ice Age maxima (in the mid-1800s), Mont Miné glacier retreated approximately by 2.53 km in length and 130m in height as of 2019 (Curry et al., 2005; Nicolussi et al. 2022). Moraines deposited during this deglaciation after the later advance are still visible (Figure 1A). In the period from 1961 to 1990, the annual averages of temperature in this glacier foreland were around 0-2°C and roughly increased by 2°C (0-4°C) in the period from 1991 to 2020. The glacier forelands received approximately 700– 1100 mm of mean annual precipitation in the period from 1961 to 2020 (data from https://map.geo.admin.ch).

**Figure 1.**
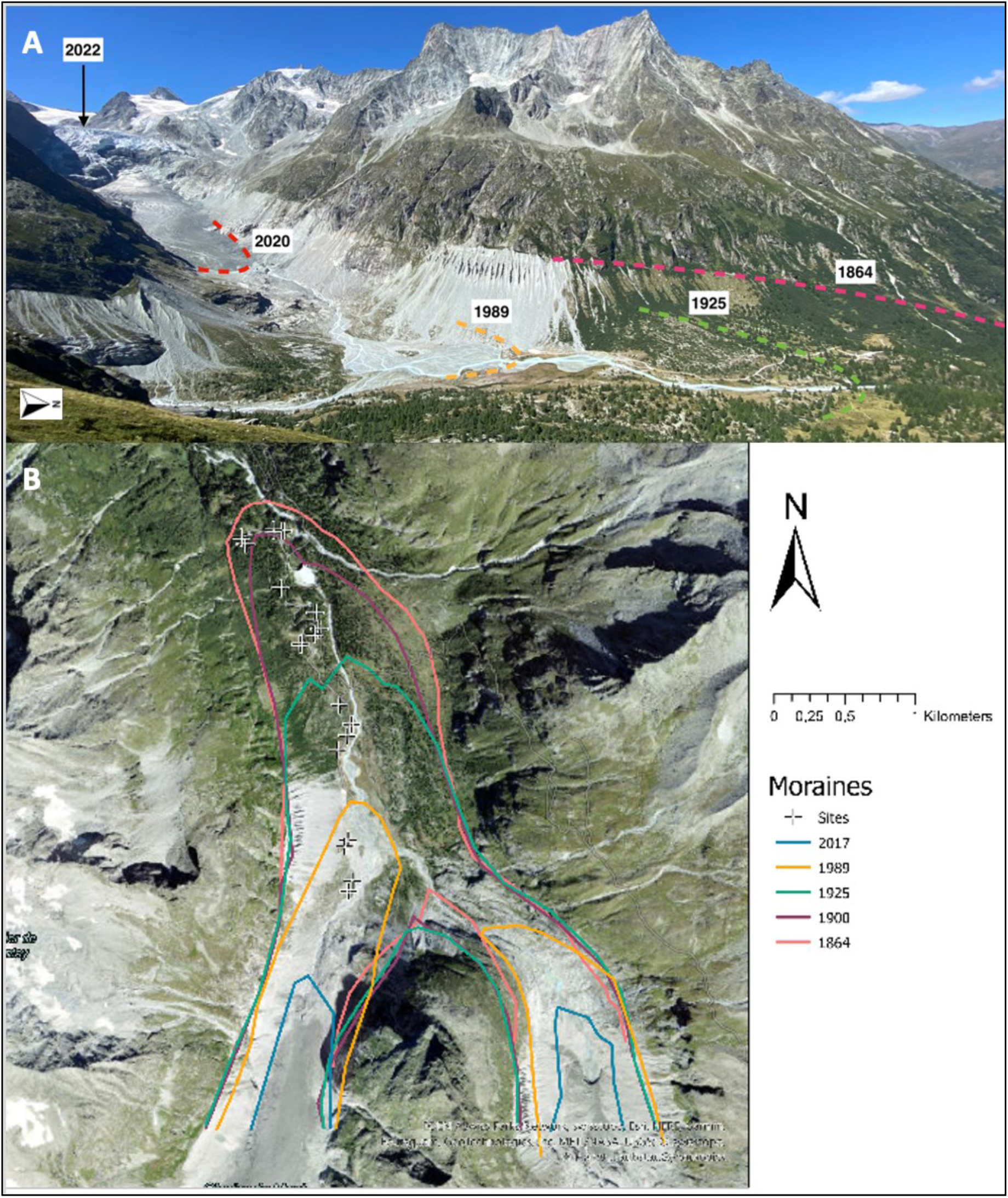
Mont Miné glacier and sampling locations. (A) View of Mont Miné glacier from the path to Bricola hut with visible moraines highlighted (photo: @B.N.Tu taken in July 2022, adapted from Nicolussi et. al. 2022). (B) Glacier foreland. Lines represent four main moraines deposit in 1864 (stage 4: pink), 1900 (stage 3: purple), 1925 (stage 2: green), 1989 (stage 1: yellow) and the present glacier (2017). Sampling sites (plots) are represented by (+) (figure: @Nora Khelidj)

Based on the existing geochronology of the Mont Miné glacier (Curry et al., 2005, Lambiel et al., 2016; Nicolussi et al., 2022) and our additional reconstruction (https://map.geo.admin.ch), we divided the Mont Miné foreland into four stages (S1, S2, S3, S4). They range from recently ice-free terrains to the end of the Little Ice Age. Stages S1, S2, S3 and S4 represent terrain deglaciated since 1989, 1925, 1900 and 1864, respectively (Figure 1) (Curry et al., 2005, Lambiel et al., 2016; Nicolussi et al., 2022). We defined the age (x) of each stage (i) equivalent to the average years since glacier retreat between two moraines as 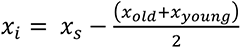 (Losapio et al., 2021a), where x_s_ is the year of sampling, x_old_ and x_young_ are the geochronological information (year) of older and younger moraines next to communities, respectively. Average ages resulted as 16.5 years for stage 1, 65 years for stage 2, 109.5 years for stage 3 and 140 years for stage 4. Stage S1 is the part of the foreland which lies in the uppermost (most recently deglaciated) sector, and S4 comprises the furthest part of the foreland from the present-day ice margins (Figure 1). We set up four plots in each of those different stages, for a total of 16 plots to carry out surveys of biodiversity to investigate ecological network dynamics. Each of the plot was 3x3 m (9m^2^) (Figure 1).

### 2. Sampling

The survey was carried out during the flowering seasons (from mid-June to the end of July) in 2022. All plots were chosen on one side of the river where the elevation gradient (from 1961 to 2000 m a.s.l) does not change much to minimize the effect of altitude on our study and is also considered to cover most of the typical vegetation at each stage as it was not disturbed by human construction activities as the side over Ferpècle glacier (Figure 2).

**Figure 2.**
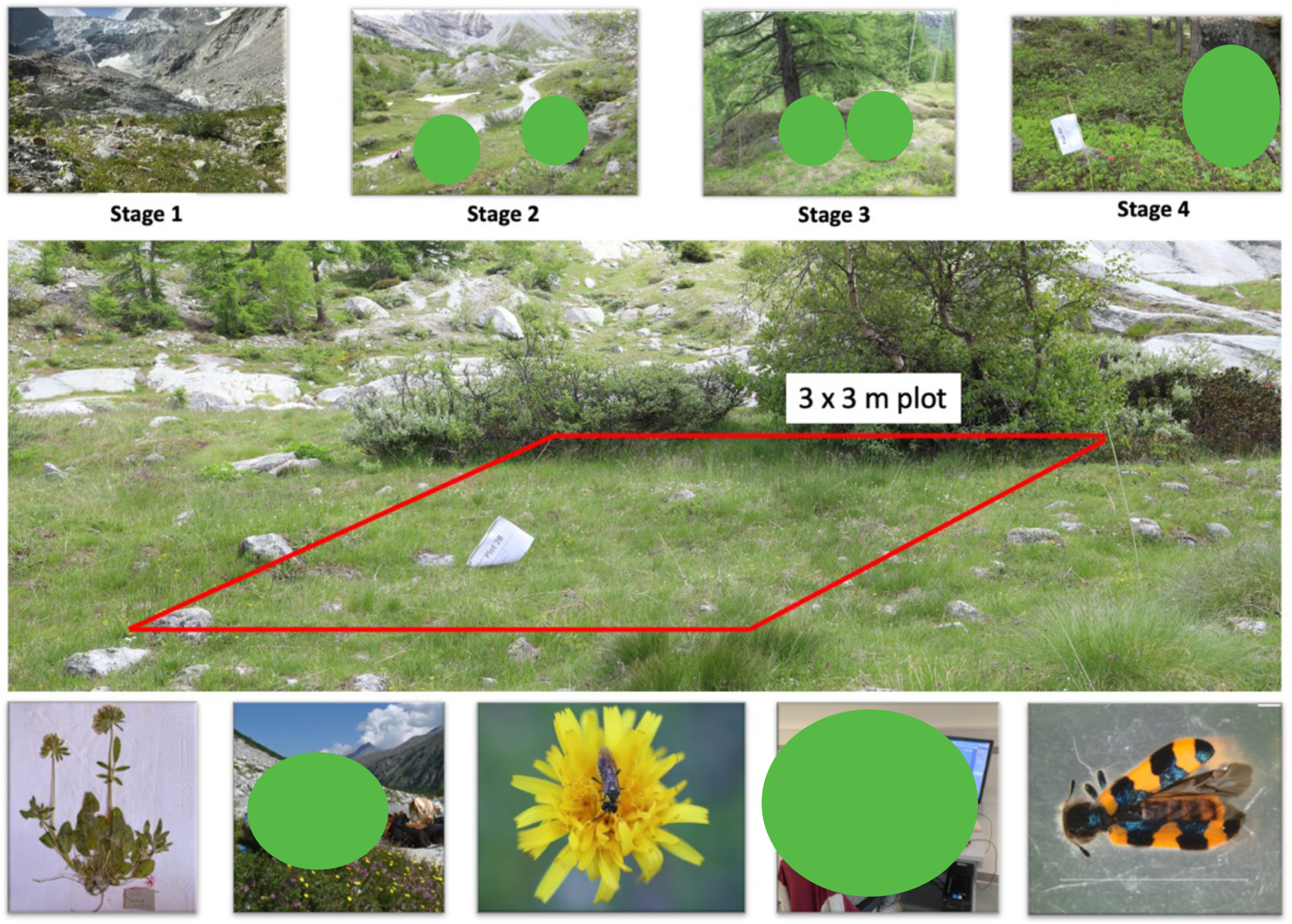
Study area in Mont Miné glacier foreland. Four stage, plot size and sampling activities are shown (Methods for details) (Photo: T.B.N, Maëlle Cornut)

#### Vegetation sampling

The vegetation survey was performed inside plots and their 1 m buffer zone. We identified all plant species (using Flora Helvetica and an updated version from the website https://www.infoflora.ch) and recorded species coverage percentage and flowering time. For flowering data, we conducted 4 replicates over time, about 15 days apart each time (Figure 2).

#### Insect sampling

Plant–insect interactions were studied by sampling insects on plants. All insects visiting flowers or staying in vegetation were sampled by an entomological aspirator, Falcon tube, sweep net or by observation and recording. We used two complementary sampling methods: quadrat and transect (Gibson et al., 2011; Martínez-Núnez et al., 2021; Grange et al., 2021). Quadrat consisted in collecting and recording insects which were on plants (flowers, leaves, branches, etc.) and on the ground inside plots and a 1m buffer zone. We separated insects caught on flowers by plant species, and those caught on leaves, branches and ground were gathered in the vegetation group. Transect consisted in collecting and recording insects which are visiting flowers on two lines (each line was 25m long and 1 m wide, 50m^2^ in total) across the plot and by assigning visitor insects to each visited plant species.

We conducted insect surveys from 9 am to 5 pm on sunny and low-wind days (Fijen et al., 2018; Martínez-Núnez et al., 2021). We monitored 30 minutes for each sampling method and did 4 replicates for quadrat (64 samples in total) and 3 replicates for transect (48 samples in total). Quadrat and transect were performed alternately and the order of plots in each replicate was chosen randomly to minimize the influence of sampling time on insect activity. Collected insects were preserved in ethanol 90 % and then stored at -20°C. Insects were identified at the highest taxonomic level, if possible at species level by specialist taxonomists, otherwise at family or order level. Microscope with high resolution was used to take photos of insect specimens for identification. We created a citizen science project on iNaturalist platform (https://www.inaturalist.org/proiects/pollinator-diversity-at-ferpecle-glacier-ecosystems) to gather further taxonomic information and for public outreach.

### 3. Data analysis

At the end of the fieldwork and labwork for insect identification, we proceeded with analyzing the data by looking at biodiversity of plants and insects first and then analyzing plant–insect interaction networks. Finally, we performed selection of the best explanatory model. All statistical analyses were conducted in R, version 4.2.2 (R Core Team 2022).

#### 3.1. Biodiversity of plants and insects

##### 3.1.1. Plant diversity

To study plant diversity, we examined the changes in plant community along the glacier retreat gradient. We took into account variables describing the changes in plant community and plant–insect interactions from the standpoint of plants for each plot: (1) plant richness, (2) Shannon diversity index at species level, (3) number of plant species flowering, (4) number of plant species visited by insects.

We first calculated plant richness as the total number of plant species occurring inside the plot and its buffer (Kiester, 2013). To estimate the changes in plant richness following glacier retreat, we fitted a Generalized Linear Model (GLM) with a Generalized Poisson distribution on our data with time (years after glacier retreat) as an effect (Bolker et al., 2008; Brooks et al., 2017).

Second, we calculated plant diversity (Shannon index) which takes into account the number of species occurring in a plot (richness) and their relative abundance (evenness). Shannon index was calculated as 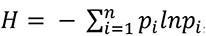, where *p_i_* is the proportional abundance of the *i*th species (*n_i_* is the number of the *i*th species) in a sample (Bersier et al., 2002; Fedor and Zvaríková 2019; Oksanene et al., 2019; Konopiński 2020). Zero is the minimum value that the Shannon diversity index can take, meaning no diversity in that habitat – there is only one species found. To estimate the effect of time (years after glacier retreat) on the changes of plant diversity, we fitted a Linear Model (LM) to our data (Chambers and Hastie, 1992).

Third, we calculated the number of plant species flowering by combining flowering data from four replicates. We considered that the changes in plant richness may affect our models, so we tested if there is any impact of plant richness by adding plant richness to the model as a predictor. To assess the combined effects (time and plant richness) on the number of plant species flowering, we fitted a Generalized Linear Mixed Model using Template Model Builder (GLMMTMB) with a Negative Binomal distribution (use for data containing a lot of zeros) and replicate was added as a random effect (Brooks et al., 2017).

Forth, we counted the number of plant species visited by insects, which means all plant species where we collected insects from both quadrat and transect data. This variable represented the relative richness of insect–pollinated plant species (Losapio et al., 2016). To estimate the combined effects (time and plant richness) on the number of plant species visited by insects, we fitted a Generalized Linear Mixed-Effects Model (GLMM) with a Poisson distribution, time and plant richness as fixed effects, method as an additional factor and replicate as a random effect on the richness of plants visited by insects. Then, we want to estimate the correlation between the number of insect-visited plant species and plant species flowering. To account for the rate of visitation in flowering plant species, we standardized by dividing plant species visited by insects for plant species flowering. To assess the effects of time on this standardization, we fitted a Linear Mixed-Effects Model via maximum likelihood (LMM) on our ratio with time as a fixed effect and replicate over time as a random effect.

##### 3.1.2. Insect diversity

To study insect diversity, we examined changes in the insect community along the glacier retreat gradient. We took into account variables describing successional changes in insect community for each plot: (1) insect richness, (2) insect abundance, and (3) insect diversity. In this study, the insect community was estimated based on orders and families because this is the highest taxonomic level that we can identify for most samples.

First, we calculated insect richness as the total number of insect families collected on flowers or vegetation in each plot for each replicate and method (n = 112). Insect abundance was calculated by the total number of individual insects collected. Insect diversity was calculated as the Shannon diversity index of insect families. To estimate the combined effects (time and plant richness) on insect richness, insect abundance and insect diversity, we fitted a GLMmTMB model with a Negative binomial distribution, method as an additional factor and replicate as a random effect on our data.

Second, to account for the response of pollinator abundance to flower density (pollinator visitation proportion), we standardized insect visitation by dividing insect abundance for flower diversity (i.e.., number of plant species flowering) at plot level (Losapio et al. 2016, 2019, 2020). To assess the effects of time on this standardization, we fitted a Linear Mixed-Effects Model via maximum likelihood (LMM) on our ratio with time as a fixed effect and replicate over time as a random effect.

#### 3.2. Plant–insect interaction networks

We created plant–insect interaction networks in each plot for each replicate and each method using the visited individual insects over each plant species (*n* = 112). To understand the structural changes of the plant–insect network along the glacier retreat gradient, for each network, we calculated interaction diversity and network complexity.

First, we calculated interaction diversity (H) using the Shannon index by computing the proportional abundance of plant–insect interactions in each plot (Bersier et al., 2002). To estimate the combined effects (time and plant richness) in plant–insect interaction diversity, we fitted a LMM with time and plant richness as fixed effects, method as an additional factor and replicate as a random effect on our data.

Second, we measured network complexity (C) by calculating the Connectance index. Connectance is the proportion of realized links from the pool of all possible interactions between the species of a network (C = links/species^2^) (Pimm, 1984; Bersier et al., 2002; Dunne et al., 2002; Heleno et al., 2012; Adedoja et al., 2018). In this study, connectance was calculated by computing the number of visiting links divided by the quantified number of plant species times the number of insect families. To estimate the effects of time in plant– insect network complexity, we fitted a LMM with time as fixed effects, method as an additional factor and replicate as a random effect on our data.

Then, to visualize interactions at local level between each family of insects and plant species, bipartite networks were constructed using the abundance of insects in a sample as a measure of relative interaction strength (Dormann et al., 2008, 2022). A bipartite network was formed from two sets of nodes (here are insect families as upper-level taxa and plant species as lower-level taxa) and connected by a set of edges. Each edge linking two different taxa belonging to different node sets presented the total interaction between them. The size of edges is proportional to the size of visitation (calculated by the number of insect individuals visiting the corresponding plant species). The size of boxes represented the size of visitor communities (the number of insect individuals for upper boxes and the number of individual plants visited by insects for lower boxes) (Dormann et al., 2008, 2022; Robinson et al., 2018; Losapio et al., 2015).

#### 3.3. Selecting the fitted model

To assess which model is the best for explaining our data, we fitted different models by increasing their complexity (Table 1). We started with a NULL model containing Intercept only. Intercept represents the mean value of the response variable when all the predictor variables in the model are equal to 0. We added a linear term for time, then followed by a quadratic term for time, and plant richness as a fixed effect. Method was added as an additional factor and replicate as a random effect (Table 1). Then, we run nested (type-I) ANOVA to assess all models and select the best-fitted model based on Deviance and AIC (Akaike information criterion). The lower values of Deviance and AIC are indication of better models (AkaikePetrov and Csaki, 1973; Portet, 2020).

**Table 1.** The formula of different model types. (Y) represents the different responses of the model (i.e., plant richness, plant diversity, insect richness, insect abundance, insect diversity, interaction diversity and network complexity); (X) represents to predictor (time and plant richness); (a) represents to Intercept; (b,c) are numerical constants

| Model type | Equation |
| --- | --- |
| Null model | $Y = a$ |
| Linear model | $Y = a + bX$ |
| Quadratic model | $Y = a + bX + cX^2$ |

Then, for each model, we used type-II ANOVA to have general information on variance explained by different predictors and we estimated model parameters to have information about the direction of effects, standard errors, and p-values. The combination of these methods allowed us to test for significant differences between single or combined effects on each response variable (Lenth, 2019). Then we presented the results with both observed data and fitted model to visualize the change in plant diversity, insect diversity and their interactions under the effect of glacier retreat (Figure 3-7).

**Figure 3.**
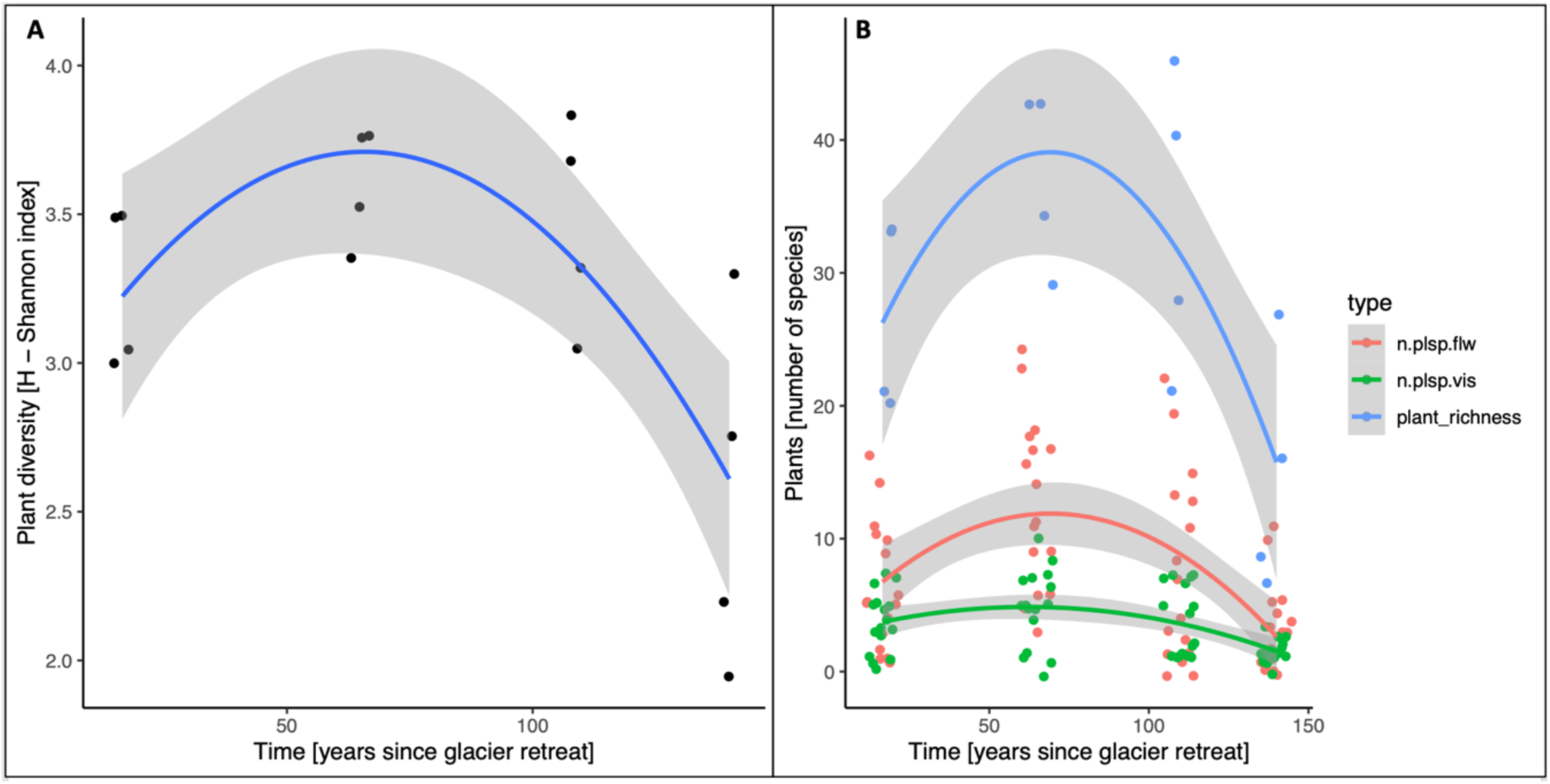
Plant community changes along the glacier retreat gradient. (A) Plant diversity (Shannon index); (B) The combined plot contained plant richness (blue), plant species visited by insects (green) and plant species flowering (red) changing along the glacier retreat gradient (calculated by number of plant species)

## RESULTS

In our study area, the total of 16 plots (of 4 stages) were set up in the summer of 2022 to follow the glacier retreat gradient for about 1.7 km length within an elevation range of 40 m a.s.l (Mont Miné glacier, Switzerland, Figure 1B).

### 1. Plant diversity dynamics under glacier retreat

We observed a total of 130 plant species belonging to 32 families across our plots and their buffer in the glacier ecosystem. The most diverse plots were plot S3D (46 species), plot S2B and plot S2C at stage 2 with 43 plant species per plot (Table S4). The poorest was plot S4A (stage 4) with only 7 plant species. There were 10 dominant plant species occurring in more than half of our plots. They are *Achillea erba-rotta*, *Anthoxanthum alpinum*, *Anthyllis vulneraria*, *Festuca rubra*, *Larix decidua*, *Lotus corniculatus* (the most dominant, occurring in 13 plots), *Peucedanum ostruthium*, *Poa alpina*, *Rhododendron ferrugineum*, *Sempervivum montanum*.

Plant diversity (Shannon index), plant richness (number of species) and plant species flowering varied along the glacier retreat gradient (Figure 3). We observed the higher diversity of plant communities in stage 2 (max H = 3.76) and stage 3 (max H = 3.82) compared with stage 1 (max H = 3.49) and stage 4 (max H = 3.29) (Figure 3A, Table S4). Here, time (years after glacier retreat) had significant negative effects on plant richness, plant diversity and plant species flowering (ANOVA, *χ*^2^ = 17.7, 2.4, 44.7, respectively, all *p* <0.05; all Estimate below 0; Figure 3; Table S1).

For the changes of plant species visited by insects, results showed the significant negative effects of time (ANOVA, *χ*^2^ = 12.9, *p* = 0.001; Estimate of linear term = -4.2, *p* = 0.02) and the significant positive effects of plant richness (ANOVA, *χ*^2^ = 20.9, *p* < 0.001; Estimate = 0.02, *p* < 0.001; Figure 4, Table S1). Instead, we observed no significant effect of the combined factors (time and plant richness) in the richness of insect-visited plant species (ANOVA, *χ*^2^ = 2.3, *p* = 0.3; Estimate = 0.08 and 0.04 in linear and quadratic model, respectively, all *p* > 0.05).

**Figure 4.**
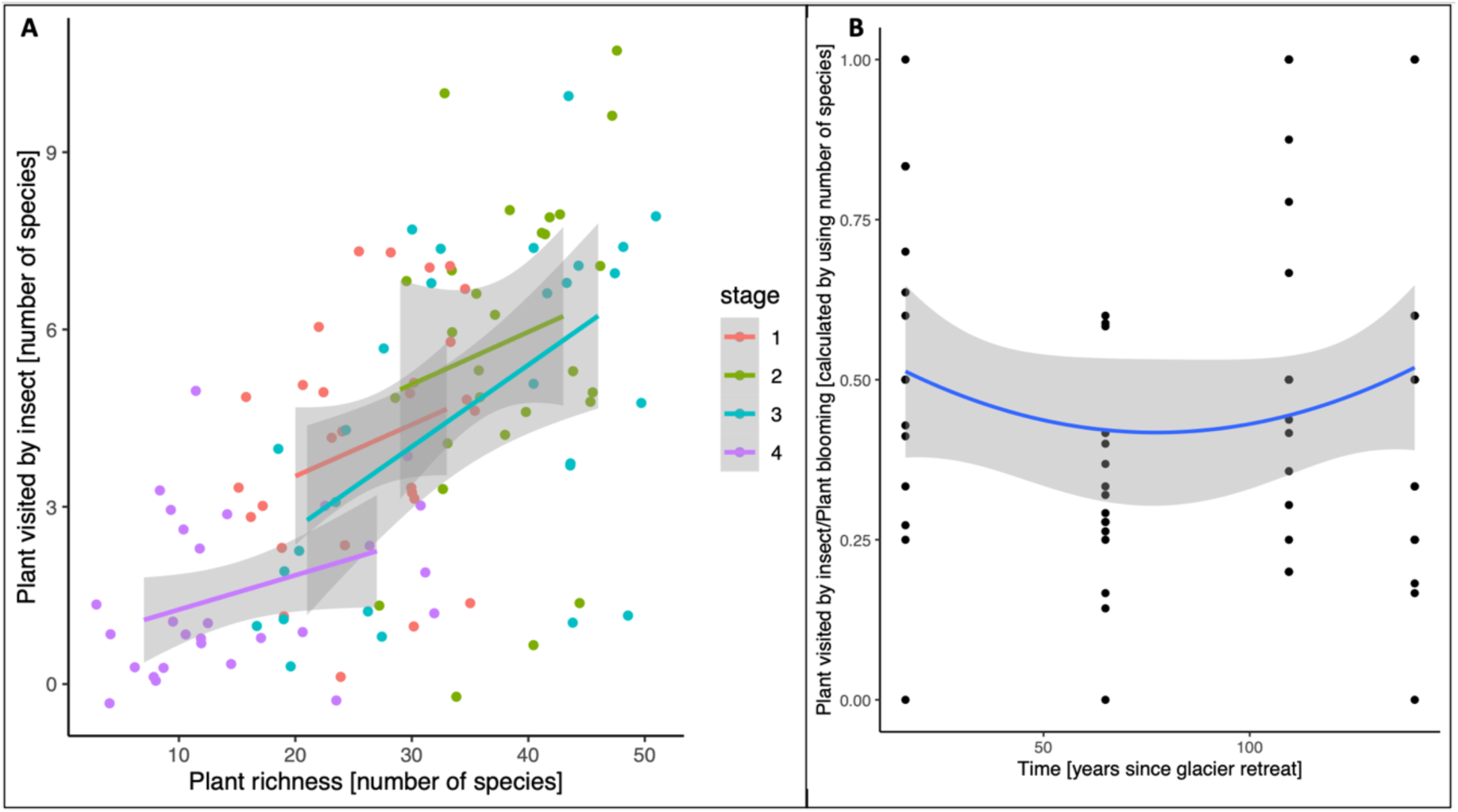
The richness of insect-visited plant species along the glacier retreat gradient. (A) The change of insect-visited plant species with positive effect added from plant richness through four stages (1, 2, 3, 4); (B) Constant model of insect-visited plant species divided by plant species flowering

We observed the constant model of plant species visited by insects divided by plant species flowering with a ratio of approximately 0.5. There were no significant differences in this ratio over time (years after glacier retreat) (ANOVA, *χ*^2^ = 1.6, *p* = 0.4; all Estimate close to zero, positive value, all *p* > 0.05; Figure 4, Table S1).

### 2. Insect diversity dynamics under glacier retreat

We collected in total of 1159 insect individuals belonging to 85 families in the quadrat data, and 923 insect individuals belonging to 50 families in the transect data (Table S4). Diptera (468 individuals from 13 families), Hymenoptera (294 individuals from 16 families), Coleoptera (169 individuals from 11 families) and Hemiptera (95 individuals from 11 families), were the most abundant orders in quadrat data. Diptera (462 individuals from 4 families), Hymenoptera (257 individuals from 8 families), Coleoptera (98 individuals from 11 families), Lepidoptera (45 individuals from 7 families), were the most abundant orders in transect data.

Insect richness, insect abundance (number of individuals) and insect diversity (Shannon index) varied along the glacier retreat gradient (Figure 5). First, for insect richness, we observed the significant negative effects of time (ANOVA, *χ*^2^ = 14.3, *p* < 0.001; Estimate in linear term = -2.8, *p* = 0.03; Figure 5A; Table S2) and the significant positive effects of plant richness (ANOVA, *χ*^2^ = 6.4, *p* = 0.01; Estimate = 0.01, *p* = 0.03; Table S2). Instead, we observed no significant effect of the combined factors (time and plant richness) in the insect richness (ANOVA, *χ*^2^ = 1, *p* = 0.5; Estimate = 0.04 and -0.002 in linear and quadratic model, respectively, all *p* > 0.05).

**Figure 5.**
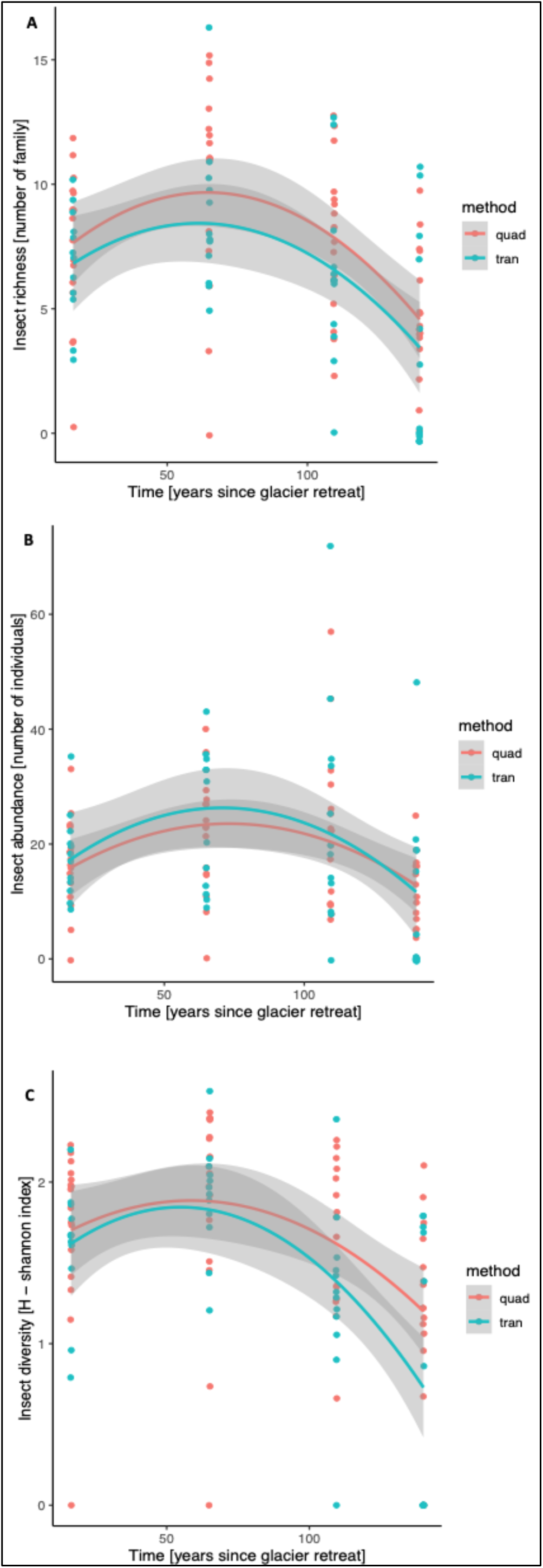
Insect community changes along the glacier retreat gradient with two collected methods (quad = quadrat method, tran = transect method). (A) Insect richness (number of insect species); (B) Insect abundance (number of insect individuals); (C) Insect diversity (Shannon index).

Second, for insect abundance, time did not affect (ANOVA, *χ*^2^ = 0.4, *p* = 0.8; Estimate in linear term = -5.3, *p* < 0.001; Figure 5B; Table S2) but we observed the significant positive effects of plant richness (ANOVA, *χ*^2^ = 15, *p* < 0.001; Estimate = 0.02, *p* = 0.003; Table S2). Additionally, the combined effects (time and plant richness) had significant positive effects in insect abundance (ANOVA, *χ*^2^ = 6.3, *p* = 0.04; Estimate in linear model = 0.19, *p* = 0.01; Table S2).

Third, for insect diversity, in quadrat data, we observed the highest average diversity of insect communities in stage 2 (max H = 2.01) and lowest in stage 4 (min H = 0.73); in transect data, highest average diversity in stage 2 (max H = 2.09) and lowest in stage 4 (min H = 0.46) (Table S4). We observed the significant negative effects of time (ANOVA, *χ*^2^ = 15, *p* < 0.001; all Estimate below 0, all *p* > 0.05; Figure 5C; Table S2) and the significant positive effects of plant richness (ANOVA, *χ*^2^ = 5.2, *p* = 0.02; Estimate = 0.01, *p* = 0.03; Table S2). However, the combined effects (time and plant richness) did not affect insect diversity (ANOVA, *χ*^2^ = 0.01, *p* = 0.9; Estimate = -0.002, 0.008 in linear and quadratic model, respectively, all *p* > 0.05; Table S2).

Besides, for the standardization of insect visitation by dividing insect abundance for plant species flowering at plot level, time did not affect this ratio (ANOVA, *χ*^2^ = 1.4, *p* = 0.4; all Estimate > 0, all *p* > 0.05; Table S2). There were about 5 insect individuals visiting 1 plant species which was flowering. The mostly constant model was shown in Figure 6.

**Figure 6.**
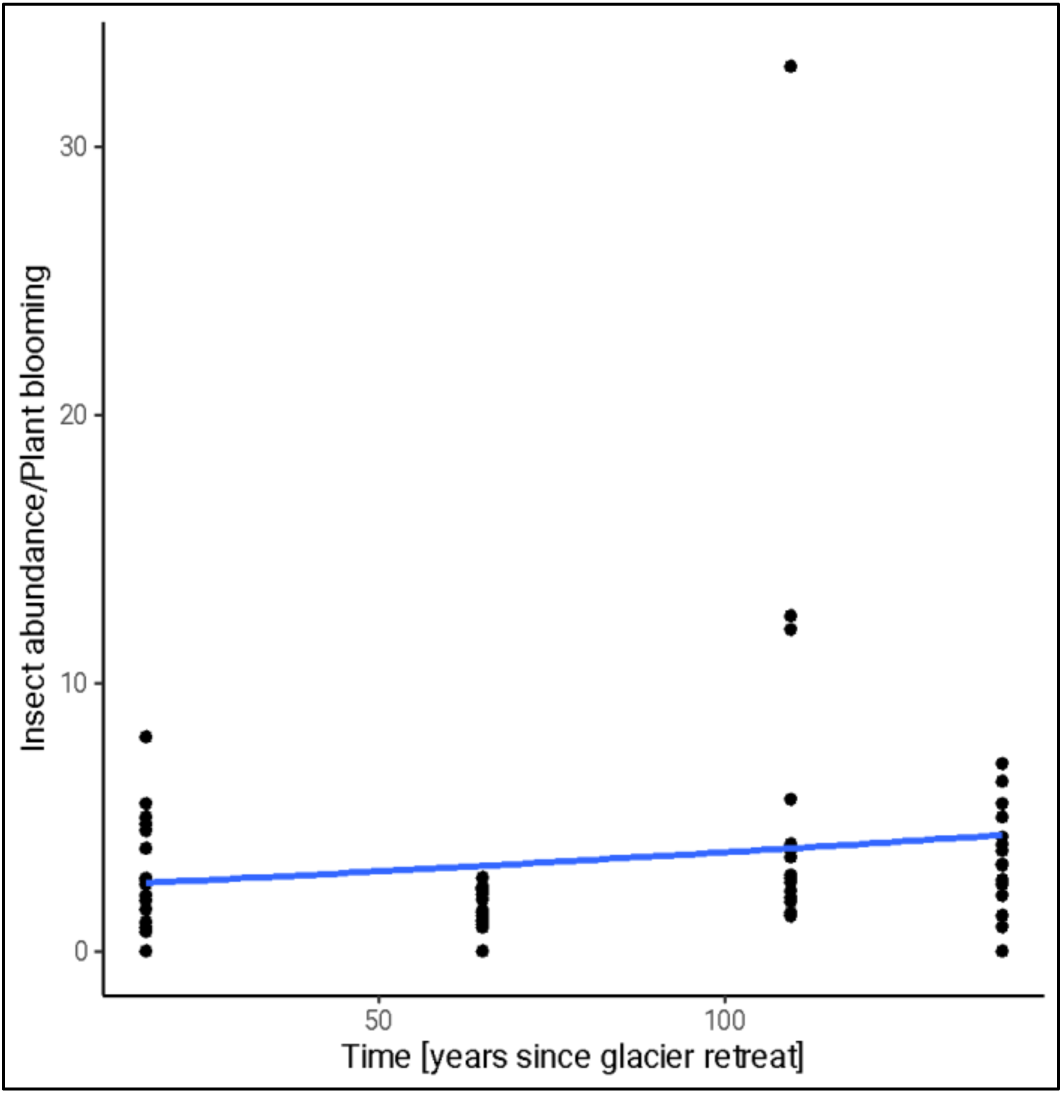
The response of pollinator abundance to flower density along the glacier retreat gradient. We standardized insect visitation by dividing insect abundance for number of plant species flowering at plot level

**Figure 7.**
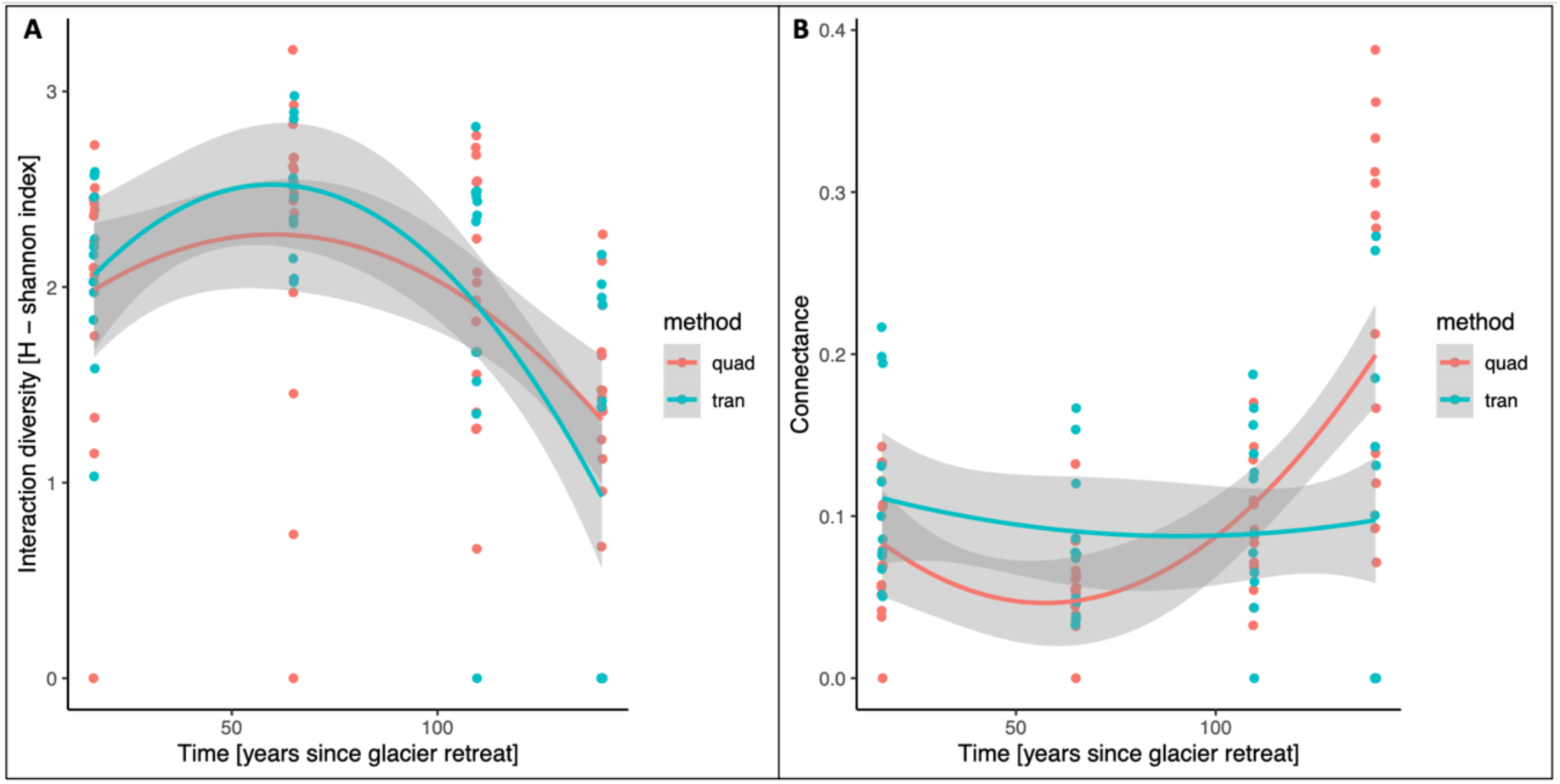
Plant–insect interaction dynamics along the glacier retreat gradient. (A) Interaction diversity (Shannon index); (B) Network complexity (connectance)

### 3. Plant–insect interaction network dynamics under glacier retreat

#### 3.1. Plant–insect interaction diversity and network complexity

We observed the changes on plant–insect interaction diversity along the glacier retreat gradient by the average Shannon index. In quadrat data, the highest interaction diversity was in stage 2 (max H = 2.36) and lowest in stage 4 (min H = 0.86). In transect data, the highest interaction diversity was in stage 2 (max H = 2.69) and lowest in stage 4 (min H = 0.46) (Table S4). We observed the significant negative effects of time (ANOVA, *χ*^2^ = 16.8, *p* < 0.001; Estimate in linear term = -3.6, *p* = 0.03; Figure 7A; Table S3) and the significant positive effects of plant richness (ANOVA, *χ*^2^ = 19.1, *p* < 0.001; Estimate = 0.02, *p* < 0.001; Table S3). Instead, there were no significant effects of the combined factors (time and plant richness) in plant–insect interaction diversity (ANOVA, *χ*^2^ = 0.7, *p* = 0.6; Estimate = 0.04 and -0.03 in linear and quadratic model, respectively, all *p* > 0.05; Table S3).

For network complexity, in quadrat data, the highest connectance index was in stage 4 (max C = 0.29) and lowest in stage 2 (min C = 0.03). In transect data, the highest connectance index was in stage 1 (max C = 0.17) and lowest in stage 2 (min C = 0.04). We observed the significant positive effects of time in connectance (ANOVA, *χ*^2^ = 22, *p* < 0.001; all Estimate > 0, all *p* < 0.001; Figure 7B; Table S3).

#### 3.2. Plant–insect interaction networks

Table 2 summarizes the data we obtained when calculating plant–insect interaction networks for four stages (1, 2, 3, 4) and separated by two different methods (quadrat and transect). For quadrat data, we observed a total of 1159 plant–insect interactions with the highest in stage 3 (366) and the lowest in stage 4 (187). For transect data, we observed a total of 932 plant–insect interactions with the highest in stage 3 (292) and the lowest in stage 4 (126).

**Table 2.** Number of insect and plant taxa in their interaction networks counted for four stages (1, 2, 3, 4) and two methods (quadrat and transect). Total = the total number of observed groups; order = number of identified insect orders; family = number of identified insect families; no id. = number of unidentified insects; species = number of identified plant species

| Method | Quadrat |  |  |  | Transect |  |  |  |
| --- | --- | --- | --- | --- | --- | --- | --- | --- |
| Stage | 1 | 2 | 3 | 4 | 1 | 2 | 3 | 4 |
|  | (fig.7A) | (fig.7B) | (fig.7C) | (fig.7D) | (fig.7E) | (fig.7F) | (fig.7G) | (fig.7H) |
| **Insects** |  |  |  |  |  |  |  |  |
| _total | 38 | 52 | 47 | 26 | 27 | 33 | 29 | 19 |
| _order | 7 | 11 | 10 | 8 | 6 | 10 | 8 | 6 |
| _family | 34 | 48 | 41 | 23 | 23 | 29 | 26 | 16 |
| _no id. | 1 | 1 | 1 | 1 | 1 | 1 |  |  |
| **Plants** |  |  |  |  |  |  |  |  |
| _total | 11 | 35 | 24 | 3 | 23 | 41 | 33 | 12 |
| _species | 10 | 34 | 23 | 2 | 23 | 41 | 33 | 12 |
| _vegetation | 1 | 1 | 1 | 1 |  |  |  |  |
| **Number of interactions** | 263 | 343 | 366 | 187 | 215 | 290 | 292 | 126 |

Then, we visualized the plant–insect interaction via bipartite networks (Figure 8). The interaction diversity and network complexity can be represented through the set of edges with each edge linking two different taxa belonging to different node sets.

**Figure 8.**
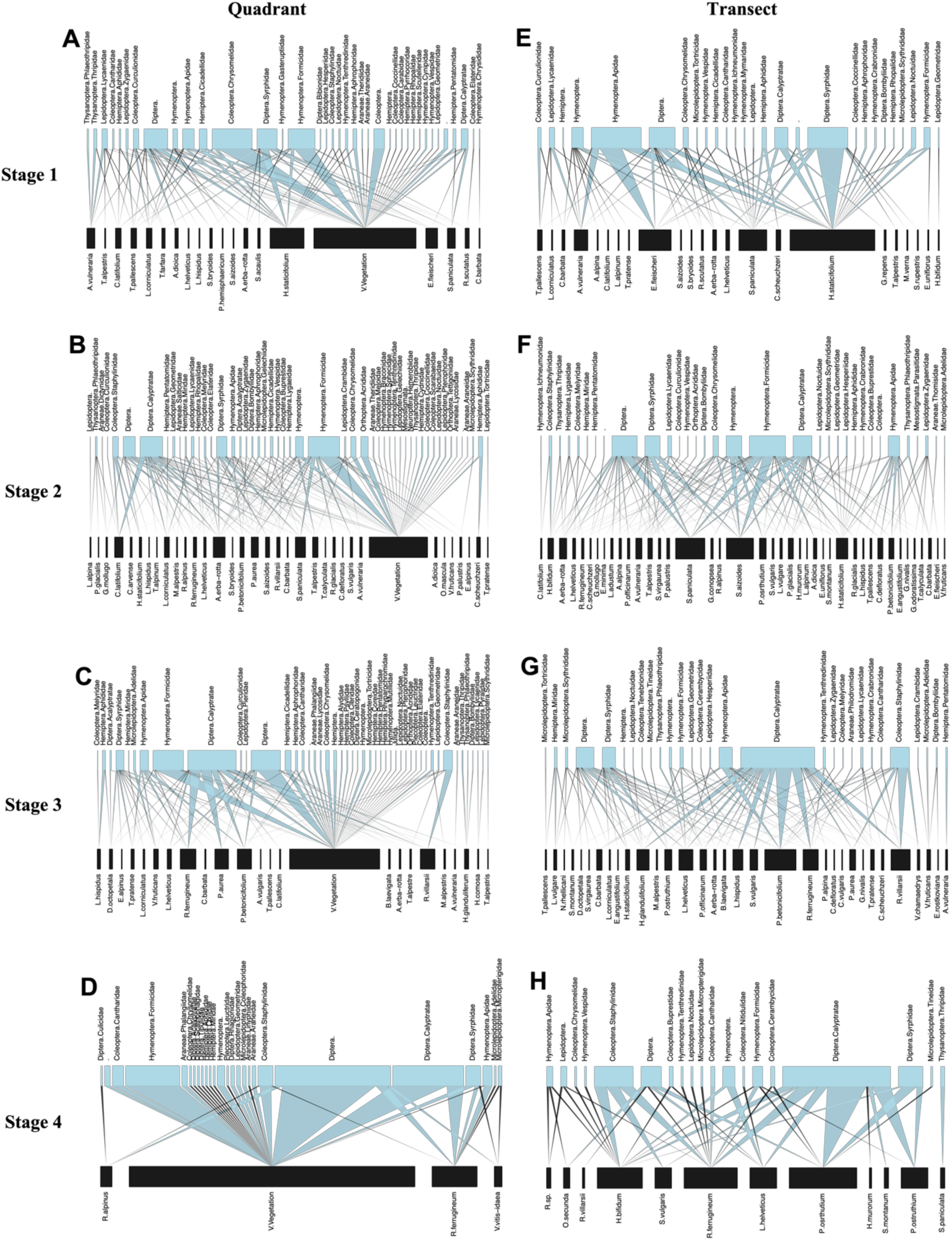
Plant–insect interaction networks for four stages along the glacier retreat gradient (Upper boxes: order and family name of insects, Lower boxes: species name of plants; Number of interactions (int): black edge: int = 1, blue edge: int > 1; A, B, C, D: networks built from quadrat data; E, F, G, H: networks built from transect data)

For the networks built from quadrat data, we observed in stage 1 that the most active pollinators were Diptera, followed by Hymenoptera (especially family Apidae) and the most attractive flower was *Hieracium staticifolium* (Figure 8A). In stage 2 and stage 3, Diptera still was the most abundant pollinators while the other groups are relatively equal, and flowering species were quite diverse (Figure 8B, 8C). In stage 4, with only 3 flowering species, Diptera still was the most abundant group and followed by Hymenoptera, but most of the interactions were on vegetation (Figure 8D).

For the networks built from transect data which was only focused on pollination, in stage 1, we observed an equal interaction of Hymenoptera (Apidae) and Diptera on flowers, while Diptera focused more on *Hieracium staticifolium*, Apidae was attracted more by *Epilobium fleischeri* (Figure 8E). In stage 2, Diptera was the most abundant insect in the network and then following by Hymenoptera (Formicidae) with more diverse flowering species (Figure 8F). In stage 3, we observed an increase in the abundance of Diptera, then following by Coleoptera (Staphylinidae) (Figure 8G). In stage 4, Diptera and Coleoptera (Staphylinidae) still were the most abundant insects in the network, while Diptera was attracted by *Peucedanum ostruthium*, Staphylinidae was found more on *Hieracium bifidum*, *Silene vulgaris* and *Rhododendron ferrugineum* (Figure 8H).

## DISCUSSION

As in several previous studies, the foreland created after glacier retreat was an extremely suitable area for studying ecological succession, especially in the context of rapid and harsh global warming (Erschbamer and Caccianiga, 2016; Ficetola et al., 2021; Rosero et al., 2021). In Mont Miné glacier foreland, we could see the sequence of all the stages of ecological succession which were relevant with 4 main stages of 140 years of deglaciation since 1864 (Figure 1). The new areas in front of the glacier were still devoid of plants, while in stage 1, pioneer species established and created flower-rich habitats. Following intermediate stages, the woody plants occurred and become more abundant in the late succession. Some adaptative species had advantages, developed stronger and become more common while some others dwindled or may lead to extinction. We observed the change in plant and insect communities and also their interaction network under glacier retreat in our study area.

### 1. Biodiversity dynamics under glacier retreat

#### 1.1. The decrease of plant diversity

In total, we identified 130 plant species of 32 families within our study area. Among those, only six plant species such as *Anthoxanthum alpinum*, *Festuca rubra*, *Larix decidua*, *Lotus corniculatus*, *Poa alpina*, *Rhododendron ferrugineum* occurred in all four stages. At family level, we observed Asteraceae, Poaceae, Fabaceae, Ericaceae as the most abundant families in the study area. In real-world ecosystems, the diversity Shannon index range (H) is usually from 1.5 to 3.5, so the plant communities in Mont Miné glacier foreland were at quite high diversity (average H = 3.32, highest H = 3.82, Table S4).

Our study indicated that global warming, especially glacier retreat in this case affected plant diversity. Our model predictions showed that plant diversity (plant species and Shannon index) and the number of plant species flowering rapidly increased at the beginning of glacier retreat (ca. 10 years after glacier retreat). However, glacier retreat had negative effects on plant communities by declining both plant richness, plant diversity and number of plant species flowering once glacier extinction (ca. 60 years after glacier retreat) (Figure 3). In fact, the initial increase of plant diversity can be explained by the appearance of pioneers from primary succession, which improved the barren soils and facilitated newcomers. The peak of plant diversity here (ca. 60 years after glacier retreat) represented the ability of species richness to colonize the free area in the early stage after deglaciation when the environmental conditions become favourable. These results align with previous studies showing the increase of plant diversity just after the glacier retreat (Frenot et al., 1998; Inouye, 2020; Losapio et al., 2021a). However, not the whole communities can gain benefits forever. We found that pioneers and some early plant species (such as *Cerastium arvense, Cerastium latifolium, Epilobium fleischeri, Linaria alpina, Oxyria digyna, Phleum alpinum, Saxifraga aizoides, Tussilago farfara*) decreased and disappeared at stage 4 (ca. 140 years after glacier retreat) (Table S5). Those pioneers were estimated to face extinction around 100 years after glacier retreat at some former researches (Caccianiga et al., 2006; Erschbamer, 2007; Losapio et al., 2021a). We also observed the sharing of plant species occurrence from these middle-aged plots with other plots in their study area (Table S5). That means there is a replacement of plant pioneer species by others. This evidence supported the majority of successional patterns of the plant communities was “replacement”, emphasized a major role of competition in stable habitat and can explain the decrease in diversity (Ficetola et al., 2021).

In the study on the effects of glacier retreat in plant primary succession 2-13 years after glacier retreat, one found a rapid decrease in both plant population level and the community levels (species richness and abundance) just 13 years after glacier retreat (from 21 species to 13 species) (Anthelme et al., 2021). Our results were more consistent with the study of plant diversity in succession (Caccianiga et al., 2006), multi-diversity in Ödenwinkel glacier (Junker et al., 2020), multi-taxa colonization (Rosero et al., 2021) indicating that the decline of plant diversity started within 100 years after glacier retreat. However, the rate of colonization and replacement might depend on local conditions (such as micro-environment, other abiotic factors, etc.,). For instance, in the harsh area with an average temperature lower, the successional rate can be slow while in the area with higher temperatures (i.e., tropical glaciers) this rate might be much higher (Franzén et al., 2019; Anthelme et al., 2021, 2022). Furthermore, this confirmed the fact that time (years after glacier retreat) had strong negative effects on plant diversity (Erschbamer and Caccianiga, 2016; Ficetola et al., 2021).

Here in this study, we use time (years after glacier retreat) as a major measurable predictor for plant diversity dynamics, but the big story behind should contain the combination effects of abiotic (such as temperature, light, soil nutrient, etc.,) and biotic factors (such as plant–plant interactions, microbiota, etc.,) (Ficetola et al., 2021).

#### 1.2. The decrease of insect diversity

Our study provided additional evidence for the effects of glacier retreat in insect communities, taking into account their richness, abundance and diversity.

We observed the highest diversity of insects (family richness and Shannon index) in stage 2 and the lowest in stage 4 (Table S4). Our model indicated that glacier retreat had negative effects on both number of insect families (richness) and insect diversity (Shannon index). Both indices increase at the early stage of deglaciation, then started declining after approximately 60 years (Figure 5). This is somehow similar to the diversity of plant communities (see the plant diversity results, Figure 3). However, the rate of insects’ decline seemed slightly slower than in plants. This is because plants are the base components of the ecosystem, especially for pollination or food web, changes in plant communities can directly affect higher trophic levels, here are insects (such as pollinators and herbivores), and then affect their interactions. Thus, it is clear observe the response of insect communities under glacier retreat is generally mediated by plant diversity (Vitasse et al., 2021). The effects of plant richness on insects’ diversity will be discussed in the later part. So, our results gave more evidence about the decrease of insect communities in mountain habitats under glacier retreat which were found in previous studies, such as the decline of terrestrial insect abundance in the land (ca. 9%) per decade in North America and some European regions (Gobbi and Lencioni, 2021; Gobbi et al., 2021).

Otherwise, for insect abundance, our results showed that time after glacier retreat did not have significant effects (Table S2). It seems that the number of insect individuals was still maintained relatively stable following glacier retreat gradient while the insect family’s composition decrease. Additionally, we observed the positive effects of the combined effect (time and plant richness) in insect abundance which can explain for this constancy following glacier retreat (discuss more in later part). Insect population structure declined at the family level may be due to the movement of insects to find more flower sources or to areas with more suitable living conditions (i.e., warmer or colder temperatures). In fact, the most abundant order of plant visitors was Diptera in both quadrat and transect data with nearly half of the total samples, followed by Hymenoptera and Coleoptera (Table S4). This major abundance of Diptera might be explained by the elevation of our study area which is from 1961 to 2000 m a.s.l. Previous studies found that at the area below 1640 m a.s.l., bees were the most dominant visitors (Adedoja et al., 2018) while there was a global shift from bees to flies as the dominant flower visitors in higher elevation systems (1922 to 3600 m a.s.l.) due to the change of air temperature (McCabe and Cobb, 2021), which is relevant with our observation. This is also consistent with the study about the dominance of flies as flower visitors in the Alps high-altitude (Lefebvre et al., 2018). The same trend was observed in the study of Bonelli et al. with added information about the effects of temperature and time (hour of the day) in the flowering visitation of insects while wind speed and the number of flowers per plant did not influence (Bonelli et al., 2022).

### 2. Plant–insect interaction network dynamics under glacier retreat

As the discussion above, plants and insects are both involved in the networks, so if all of them were influenced negatively by glacier retreat, their interactions might be affected in the same way (Vitasse et al., 2021). In fact, we observed the dynamics of plant–insect interaction diversity, initially increased at the early stage and then rapidly reduced with the glacier extinction. In total, our results indicated that glacier retreat had negative effects on interaction diversity (Shannon index, Figure 7, Table S3). It means that as the plant and insect communities decline, their interactions were also reduced following glacier retreat gradients. Otherwise, we observed the shift in plant–insect interaction networks in quadrat data, from the pollination dominant (stage 1, 2, 3) to food web dominant (stage 4) following glacier retreat (Figure 8A-D). This result agrees with the study of interactions between two plant species and their visitors under the effects of glacier retreat (Losapio et al. 2015). Thus, we hypothesize that this shift may increase the robustness of the plant–insect interaction network. Also, insects can maximize plant resources while number of plant species in general and number of flowering plant species in particular decline under glacier retreat. This process partly reflects the adaptation of insects to changes in the environment.

Plant–insect interaction networks are represented by associations (links) between species (nodes) (Figure 8) and connectance (C) was used to describe the network complexity with consequences for community stability, corresponding to the number of actual links over the number of possible links (Bersier et al., 2002; Dunne et al., 2002a, b). Any loss of species in networks can cause a change in connectance, then impact and lead to the loss of robustness (Gilbert, 2009). The low connectance networks are the most accessible ones to attack by the loss of species or by abiotic factors (Dunne et al., 2002a, b). In another way, within the interaction network, increase connectance will help to reduce the extinction of species in the network (Blüthgen et al., 2006). Our results showed that the plant–insect interaction networks in stage 2 had the lowest connectance in both quadrat and transect data. It reflected the instability of plant and insect communities in stage 2, even though there was the highest diversity of plants and insects here but these communities can be easily affected by abiotic and biotic factors (Table S3). Indeed, our results about the decrease of biodiversity under glacier retreat confirmed that. In quadrat data, we observed the highest connectance in stage 4 where pollination networks dominant shift to food web dominant, representing the stability of the ecosystem (Table S3). Transect data was focused only on flowering visitors so we found the highest connectance in stage 1. In addition, the positive effects of time (years after glacier retreat) in connectance supported the stability of networks in stage 4 which is in distance from the recent glacier. Our results were similar to the study of Adedoja et al., they observed the decline of connectance value while network diversity increased along elevations (Adedoja et al., 2018). Even though our study did not perform along the altitude gradients but both results confirmed the stability of interaction network of communities at lower zones (here in our study is communities further with glacier) but not communities with higher taxa diversity. This result is in contrast with the former observation about communities with rich partners and diversity links are usually more stable than the few species communities (Dunne, 2006) but confirmed again the negative effects of glacier retreat in biodiversity and ecological networks of sensitive ecosystems in high altitude mountains.

However, in our study, we analyzed the complexity of plant–insect interaction networks based on qualitative indices connectance (C) which strongly depended on networks size. To research deeper and get more information about interaction frequencies or sampling intensity, other quantitative indices should be studied such as the degree of specialization, the species-level index *d’* to analyze variation inside network and the network-level index *H_2_’* to compare different networks (Blüthgen et al., 2006).

### 3. Additional effects of plant richness on biodiversity under glacier retreat

As former studies mentioned, plants play an important role in a healthy functioning ecosystem and are a stronger predictor of network structure (Losapio et al., 2018). Here, we also found that plant richness strongly influences insect communities and their interactions. Precisely, first, we observed the significant negative effects of time and the significant positive effects of plant richness (Figure 4, Table S1) on number of plants visited by insects. That confirmed the important role of plant diversity in the ecosystem. The higher the richness, the higher percentage of flowering plant species, thereby increasing the ability of plants to attract insects, then leading to successful pollination (Inouye, 2020). Interestingly, we observed plant species were visited at a constant rate of approximately 0.5. This means for every 2 flowering plant species, there will be 1 plant species visited by insects. This rate represented how plants were visited by insects and it was constant with time following glacier retreat. In fact, this result was relevant to the previous experiment showed that the increase in plant diversity enhances the frequency and temporal stability of insect visitation (Losapio et al., 2020). This can be the result of expanding blossom cover and the occurrence of attractive flowering species, and also due to the stability of vegetation which can provide food or space for insects (Adedoja et al., 2018). This proved the sustainability of plant–insect relationships even under the influence of deglaciation. Insect visitors try to maximize the benefits of plant resources by seeking all the ways to exploit plants. That means although floral resources could be the most important factor which attracts insects, the richness of plant species, in general, is also extremely important to increase insect communities, thereby contributing to healthy diverse ecosystems.

Second, in the change of insect communities, besides the negative effects of time, our model indicated that plant richness affected positively insect richness and insect diversity (Table S2). Plant richness also added positive effects on insect abundance even though there was no effect from time (Table S2). However, the combined effects between plant richness and time did not have significant effects on insect richness and insect diversity, but had positive effects on insect abundance. This observation could explain the reason why insect diversity decreases under glacier retreat even when plant richness might influence to increase the insect communities.

Third, when assessing insect visitation rates by dividing insect abundance by the number of flowering plant species, we also did not see the effect of time after glacier retreat. On average, about 5 insect individuals were visiting 1 plant species which was flowering every half hour, and this rate did not change following glacier retreat gradient. Here we used insect abundance as in former studies and the number of plant species flowering following some previous results showed the influence of flower resources in insects (Adedoja et al., 2018; Grange et al., 2021). This result gave additional evidence for the sustainability of plant–insect relationships even under the influence of deglaciation. In addition, they found that visitation density can be mainly explained by the diversity of ecological variables of interest (i.e., flower areas), improve the importance of plant richness, especially flowering species in study plant–insect interactions (Grange et al., 2021). Finally, to improve the effect of plant richness, our model also showed the significant positive of plant richness in plant– insect interaction diversity (Shannon index) (Table S3).

There was still an argument in the global study about the response of biodiversity to glacier retreat, they suggested that diversity (taxon abundance and richness) of different taxonomic populations increase locally as glacier retreat (Cauvy-Fraunié and Dangles, 2019). There was still some loss (6-11% of their studied populations) but most of the losers were specialist species strictly adapted to glacial conditions. Here, our results were more consistent with other studies which indicated that glacier retreat will decrease biodiversity on a global scale (Dullinger et al., 2012; Losapio et al., 2021a; Hanusch et al., 2022). Although the losers that become extinct may be specialist species that depend entirely on glacial habitats, the implications could be much huger if we delve deeper into the relationships within the population, and then gradually to the community and the whole ecological system. It also proves the importance of studying interactions in ecology, especially in ecosystems sensitive to global warming such as glaciers in the mountains. Yet, after controlling for the effects of flower diversity, interaction frequency showed a regular pattern. Accordingly, the complexity of pollination networks and interaction diversity tended to change at constant rates with glacier retreat. However, our study only provides initial results about the impact of glacier retreat on biodiversity and plant–insect interactions in the Mont Miné glacier foreland. It is necessary to add the effects of abiotic factors on the models and analyze more network indices such as the degree of specialization (Blüthgen et al., 2006) or network eigenvector centrality (Losapio et al., 2021) to clarify this change and quantify network structure. In addition, further studies on functional groups of plants and insects in the interaction networks can also better explain the structural changes. Furthermore, our study only evaluated the effects of plant richness on biodiversity, it is necessary to further evaluate the feedback of insects to have more comprehensive view of this interaction.

In conclusion, our results confirmed the hypothesis that in the long-term under glacier retreat, plan and insect diversity will decrease, followed by their interaction networks. Plant richness plays an importance role in driving insect communities and their interactions. Increasing plant diversity would help to maintain the diversity of insects, not only pollinators but also other functional insect groups and build up stable networks. That means supporting the diversity of plants may therefore be a key strategy for halting the erosion of ecological networks under the negative impacts of global warming while increasing ecosystem functioning.

## Acknowledgments

This study was supported from the Swiss National Science Foundation (PZ00P3_202127). We are grateful to the Service des forêts, de la nature et du paysage of Canton Valais. We thank Daniele Sommaggio, Jürg Schmid, and Pierfilippo Cerretti for helping with insect identification.

---

**Table S1.** Summary statistics for the effects of time (years after glacier retreat), plant richness as predictors and method as additional factor on plant richness, plant diversity (Shannon index), plant species visited by insect, plant species flowering and ratio plant species visited by insect/plant species flowering. We first report the anova table with Wald test, degrees of freedom (df) and *P-*value followed by model estimates, standard error (SE), *t-* and *p*-values. Significant effects are highlighted in **bold**.

| Plant richness |  |  |  |  |
| --- | --- | --- | --- | --- |
| Anova table | Wald test |  | df | p |
| **Time** | 17.775 |  | 2 | **< 0.001** |
| Model parameters | Estimate | SE | t | p |
| Intercept (average plant richness) | 3.282 | 0.08 | 40.82 | **< 0.001** |
| **Effect of time in linear model** | -0.718 | 0.337 | -2.13 | **0.003** |
| **Effect of time in quadratic model** | -1.149 | 0.302 | -3.80 | **< 0.001** |
| Plant diversity |  |  |  |  |
| Anova table | Wald test |  | df | p |
| **Time** | 2.494 |  | 2 | **0.004** |
| Model parameters | Estimate | SE | t | p |
| Intercept (average plant diversity) | 3.218 | 0.096 | 33.254 | **< 0.001** |
| **Effect of time in linear model** | -0.878 | 0.387 | -2.27 | **0.04** |
| **Effect of time in quadratic model** | -1.312 | 0.387 | -3.391 | **0.004** |
| Plant species visited by insect |  |  |  |  |
| Anova table | Wald test |  | df | p |
| **Method** | 1.217 |  | 1 | 0.269 |
| **Time** | 12.978 |  | 2 | **0.001** |
| **Plant richness** | 20.998 |  | 1 | **< 0.001** |
| **Time : Plant richness** | 2.313 |  | 2 | 0.314 |
| Model parameters | Estimate | SE | t | p |
| Intercept (average plant visited by insect) | 0.227 | 0.332 | 0.685 | 0.493 |
| **Method** | 0.446 | 0.404 | 1.104 | 0.269 |
| **Effect of time in linear model** | -4.291 | 1.87 | -2.294 | **0.021** |
| **Effect of time ion quadratic model** | -2.703 | 2.137 | -1.265 | 0.205 |
| **Plant richness** | 0.027 | 0.006 | 4.188 | **< 0.001** |
| **Effect of time:plant richness in linear model** | 0.082 | 0.063 | 1.295 | 0.195 |
| **Effect of time:plant richness in quadratic model** | 0.04 | 0.068 | 0.597 | 0.55 |
| Plant species flowering |  |  |  |  |
| Anova table | Wald test |  | df | p |
| **Time** | 44.748 |  | 2 | **< 0.001** |
| Model parameters | Estimate | SE | t | p |
| Intercept (average plant blooming) | 1.714 | 0.352 | 4.86 | **< 0.001** |
| **Effect of time in linear model** | -2.145 | 0.618 | -3.471 | **< 0.001** |
| **Effect of time in quadratic model** | -3.487 | 0.575 | -6.062 | **< 0.001** |
| Plant species visited by insect/plant species flowering |  |  |  |  |
| Anova table | Wald test |  | df | p |
| **Time** | 1.677 |  | 2 | 0.432 |
| Model parameters | Estimate | SE | *t* | *p* |
| Intercept (average ratio) | 0.474 | 0.054 | 8.651 | **0.003** |
| **Effect of time in linear model** | 0.005 | 0.261 | 0.021 | 0.982 |
| **Effect of time in quadratic model** | 0.339 | 0.261 | 1.295 | 0.2 |

**Table S2.**
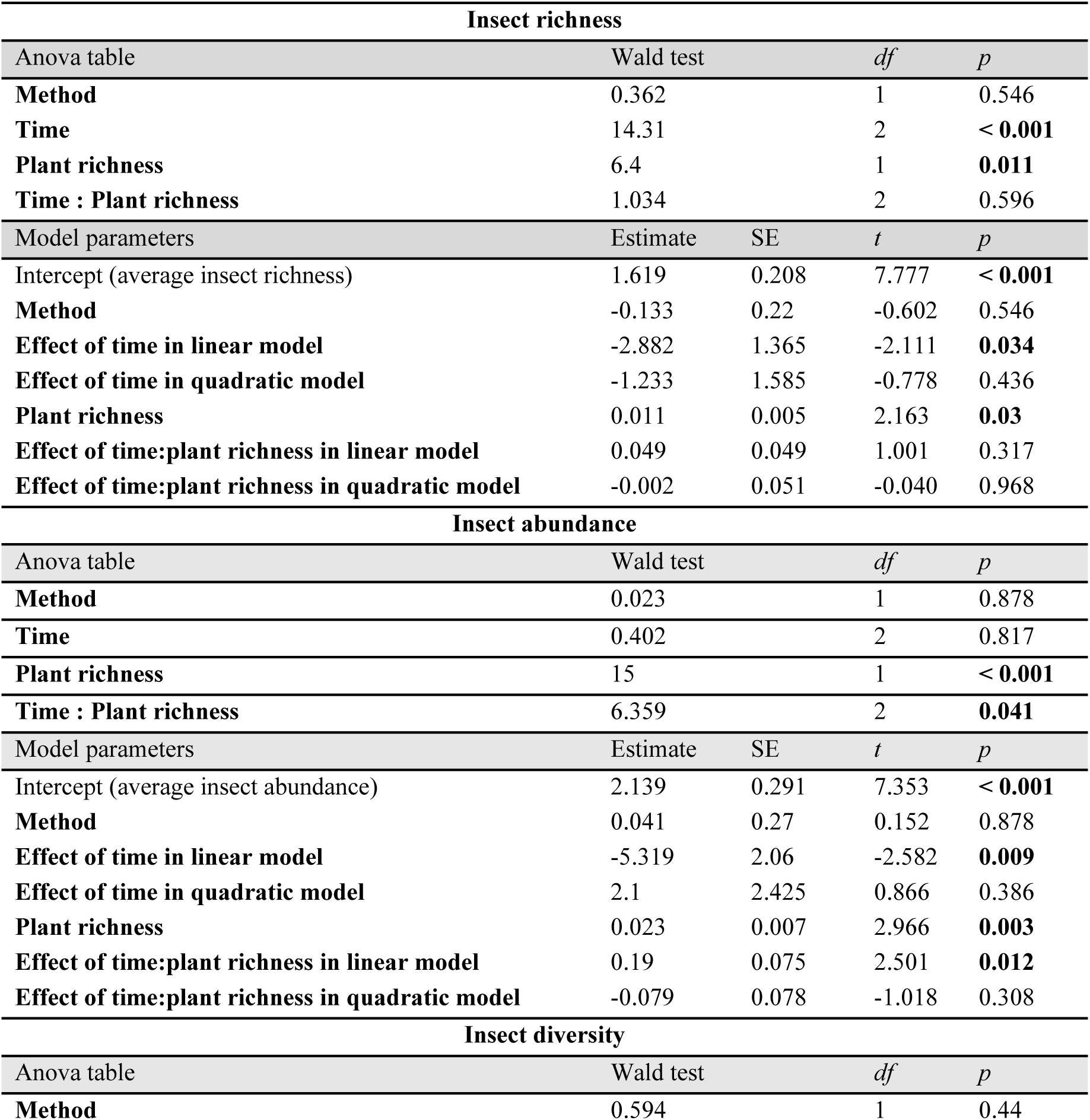
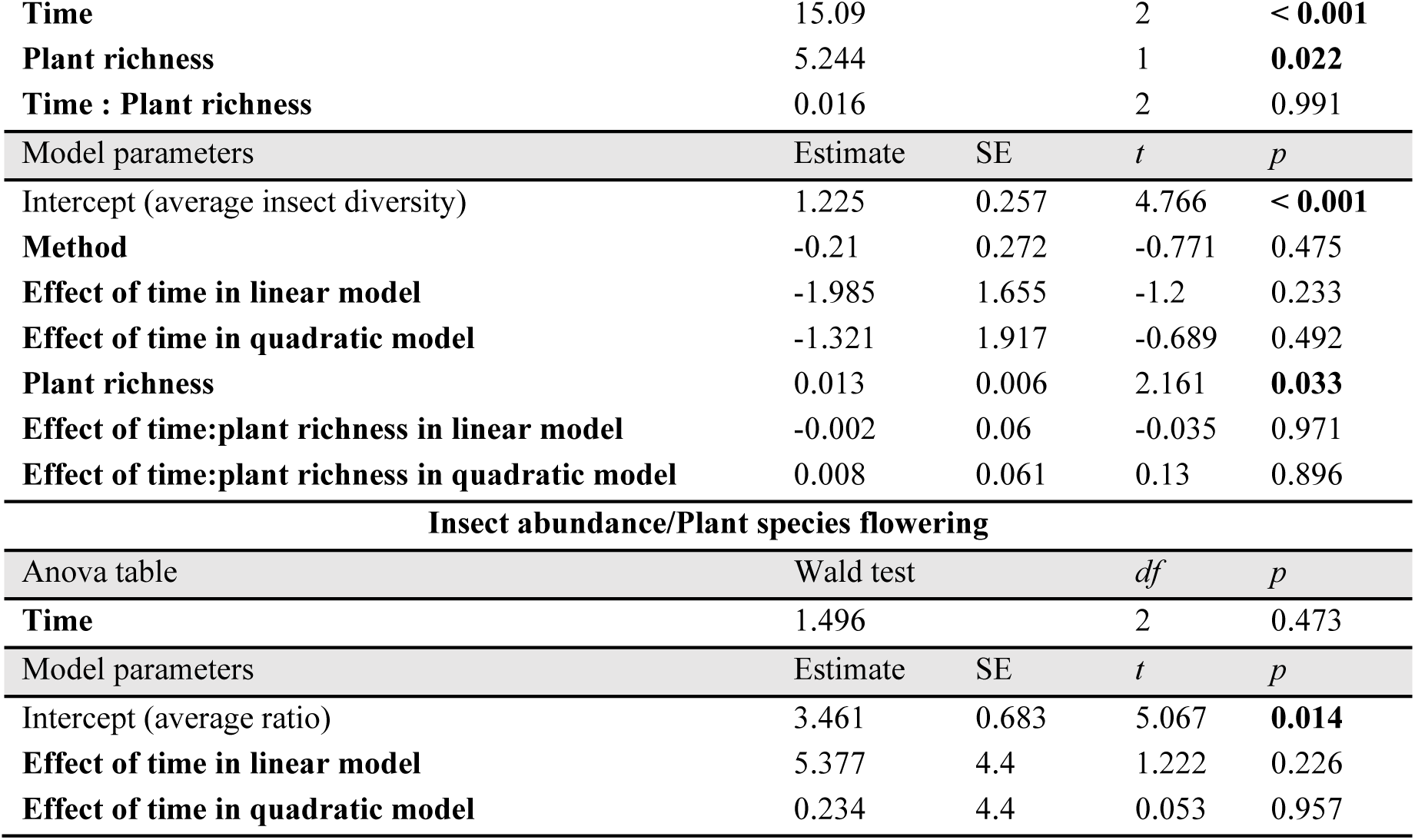
Summary statistics for the effects of time (years after glacier retreat), plant richness as predictors and method as additional factor on insect richness, insect abundance, insect diversity (Shannon index), ratio insect abundance/plant species flowering. We first report the anova table with Wald test, degrees of freedom (df) and *P-*value followed by model estimates, standard error (SE), *t-* and *P*-values. Significant effects are highlighted in **bold**.

**Table S3.** Summary statistics for the effects of time (years after glacier retreat), plant richness as predictors and method as additional factor on interaction diversity (Shannon index) and network complexity (connectance). We first report the anova table with Wald test, degrees of freedom (df) and *P-*value followed by model estimates, standard error (SE), *t-* and *P*-values. Significant effects are highlighted in **bold**.

| Interaction diversity |  |  |  |  |
| --- | --- | --- | --- | --- |
| Anova table | Wald test |  | *df* | *p* |
| **Method** | 0.001 |  | 1 | 0.969 |
| **Time** | 16.845 |  | 2 | **< 0.001** |
| **Plant richness** | 19.156 |  | 1 | **< 0.001** |
| **Time : Plant richness** | 0.715 |  | 2 | 0.699 |
| Model parameters | Estimate | SE | *t* | *p* |
| Intercept (average insect richness) | 1.148 | 0.319 | 3.593 | **0.003** |
| **Method** | -0.014 | 0.388 | -0.038 | 0.97 |
| **Effect of time in linear model** | -3.609 | 1.732 | -2.084 | **0.039** |
| **Effect of time in quadratic model** | -0.092 | 2.006 | -0.046 | 0.963 |
| **Plant richness** | 0.025 | 0.006 | 3.866 | **< 0.001** |
| **Effect of time:plant richness in linear model** | 0.049 | 0.063 | 0.771 | 0.442 |
| **Effect of time:plant richness in quadratic model** | -0.036 | 0.064 | -0.555 | 0.579 |
| Network complexity |  |  |  |  |
| Anova table | Wald test |  | *df* | *p* |
| **Method** | 0.927 |  | 1 | 0.335 |
| **Time** | 22.09 |  | 2 | **< 0.001** |
| Model parameters | Estimate | SE | *t* | *p* |
| Intercept (average insect abundance) | 0.109 | 0.01 | 10.465 | < **0.001** |
| **Method** | -0.013 | 0.014 | -0.963 | 0.337 |
| **Effect of time in linear model** | 0.232 | 0.073 | 3.18 | **0.001** |
| **Effect of time in quadratic model** | 0.252 | 0.073 | 3.461 | < **0.001** |

**Table S4.** Summary table index of plant diversity, insect diversity and their interaction networks. Plot: 4 plots (A, B, C, D) per each stage (S1, S2, S3, S4). Time: years after glacier retreat. Plant richness: n = number of plant species. Plant diversity, insect diversity, interaction diversity (Shannon index): 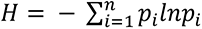. Insect abundance: n = number of individual insects. Insect richness: n = highest number of insect families observed in 4 replicates for quadrat and 3 replicates for transect. Connectance: C = links/species^2^. Method: quadrat and transect.

| Plot | S1A | S1B | S1C | S1D | S2A | S2B | S2C | S2D | S3A | S3B | S3C | S3D | S4A | S4B | S4C | S4D |
| --- | --- | --- | --- | --- | --- | --- | --- | --- | --- | --- | --- | --- | --- | --- | --- | --- |
| Time | 16.5 | 16.5 | 16.5 | 16.5 | 65 | 65 | 65 | 65 | 109.5 | 109.5 | 109.5 | 109.5 | 140 | 140 | 140 | 140 |
| **Plant community** |  |  |  |  |  |  |  |  |  |  |  |  |  |  |  |  |
| Plant richness | 20 | 21 | 33 | 33 | 29 | 43 | 43 | 34 | 40 | 21 | 28 | 46 | 7 | 9 | 16 | 27 |
| Plant diversity | 2.99 | 3.04 | 3.49 | 3.48 | 3.35 | 3.76 | 3.76 | 3.52 | 3.67 | 3.04 | 3.31 | 3.82 | 1.94 | 2.19 | 2.75 | 3.29 |
| **Insect community (quadrat data)** |  |  |  |  |  |  |  |  |  |  |  |  |  |  |  |  |
| Insect abundance | 54 | 90 | 63 | 56 | 89 | 99 | 96 | 59 | 138 | 58 | 63 | 107 | 46 | 32 | 46 | 63 |
| Insect richness | 12 | 11 | 10 | 10 | 13 | 15 | 14 | 15 | 13 | 7 | 12 | 12 | 8 | 5 | 7 | 10 |
| Insect diversity | 1.45 | 1.73 | 1.87 | 1.76 | 1.83 | 2.01 | 1.80 | 1.80 | 1.79 | 1.39 | 1.49 | 1.88 | 1.43 | 0.73 | 1.18 | 1.40 |
| **Insect community (transect data)** |  |  |  |  |  |  |  |  |  |  |  |  |  |  |  |  |
| Insect abundance | 62 | 56 | 49 | 48 | 66 | 87 | 101 | 36 | 142 | 22 | 68 | 60 | 4 | 19 | 69 | 34 |
| Insect richness | 9 | 8 | 10 | 8 | 16 | 8 | 11 | 10 | 13 | 4 | 12 | 6 | 4 | 8 | 11 | 7 |
| Insect diversity | 1.44 | 1.58 | 1.64 | 1.71 | 2.08 | 1.69 | 1.72 | 2.09 | 1.45 | 0.65 | 1.66 | 1.40 | 0.46 | 0.59 | 1.15 | 0.86 |
| **Plant–insect interactions (quadrat data)** |  |  |  |  |  |  |  |  |  |  |  |  |  |  |  |  |
| Interaction diversity | 1.78 | 2.00 | 2.06 | 2.13 | 2.23 | 2.36 | 2.28 | 2.00 | 2.34 | 1.48 | 1.83 | 2.18 | 1.53 | 0.86 | 1.18 | 1.59 |
| Connectance | 0.08 | 0.13 | 0.06 | 0.06 | 0.09 | 0.06 | 0.06 | 0.03 | 0.09 | 0.12 | 0.09 | 0.06 | 0.29 | 0.23 | 0.19 | 0.11 |
| **Plant–insect interactions (transect data)** |  |  |  |  |  |  |  |  |  |  |  |  |  |  |  |  |
| Interaction diversity | 1.93 | 1.94 | 2.12 | 2.28 | 2.50 | 2.45 | 2.69 | 2.23 | 2.59 | 1.00 | 2.11 | 2.15 | 0.46 | 0.63 | 1.37 | 1.14 |
| Connectance | 0.17 | 0.14 | 0.08 | 0.07 | 0.07 | 0.11 | 0.10 | 0.04 | 0.14 | 0.10 | 0.10 | 0.08 | 0.05 | 0.09 | 0.13 | 0.10 |

**Table S5.**
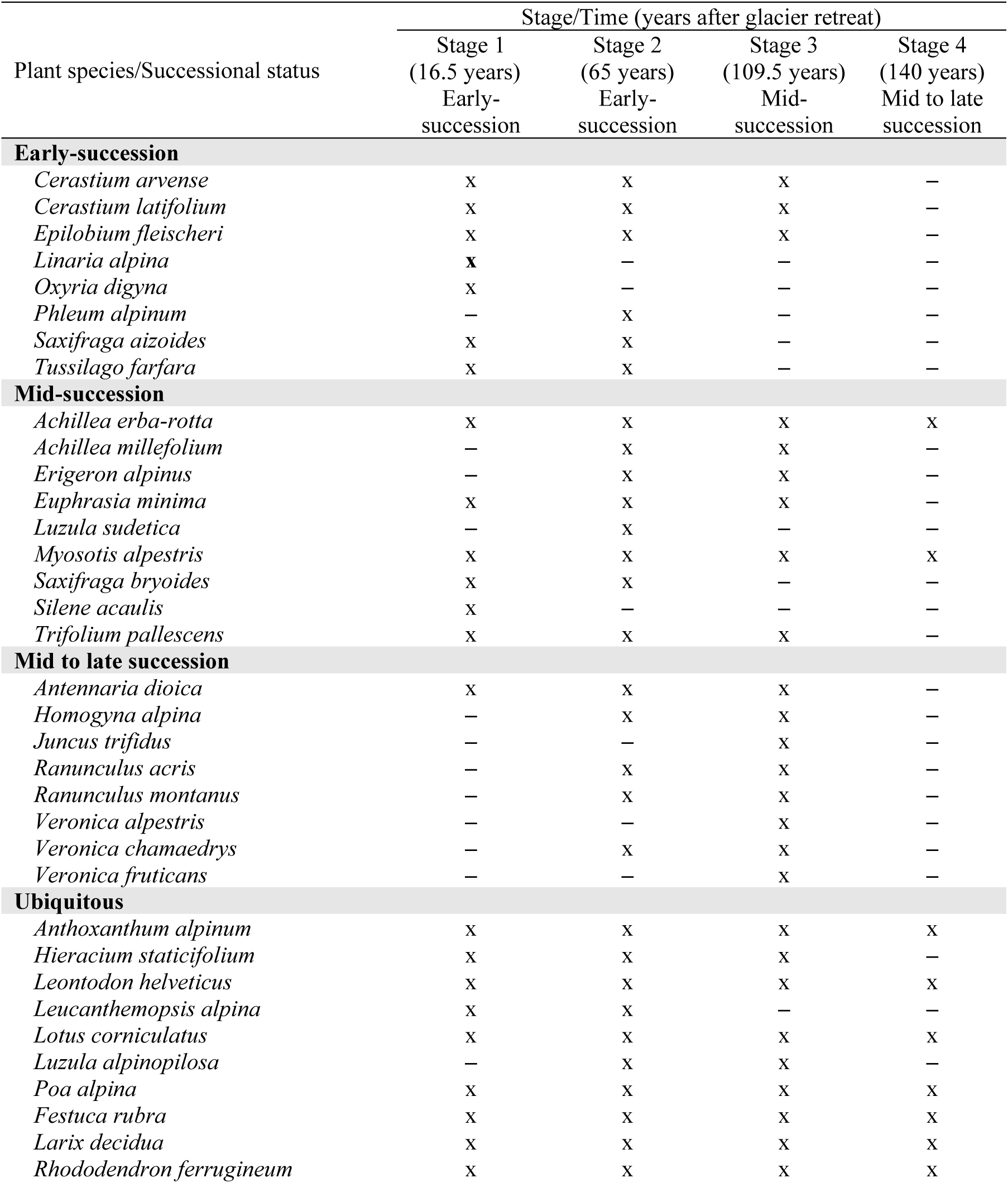
Successional status of plant species which were known occurring at different successional stages (following Caccianiga et al., 2006 and present observation of this study) in 4 stages of Mont Mine glacier forelands (x = present, **–** = absent). Plant species considered in early-succession will be not found in late succession.

## BIBLIOGRAPHY

1. Adedoja, O. A., Kehinde, T., & Samways, M. J. (2018). Insect-flower interaction networks vary among endemic pollinator taxa over an elevation gradient. PLOS ONE, 13(11), e0207453. https://doi.org/10.1371/journal.pone.0207453

2. Akaike, H. (1973) Information Theory and an Extension of the Maximum Likelihood Principle. In: Petrov, B.N. and Csaki, F., Eds., International Symposium on Information Theory, 267–281.

3. Anthelme, F., Carrasquer, I., Ceballos, J. L., & Peyre, G. (2022). Novel plant communities after glacial retreat in Colombia: (Many) losses and (few) gains. Alpine Botany, 132(2), 211–222. https://doi.org/10.1007/s00035-022-00282-1

4. Anthelme, F., Cauvy-Fraunié, S., Francou, B., Cáceres, B., & Dangles, O. (2021). Living at the Edge: Increasing Stress for Plants 2–13 Years After the Retreat of a Tropical Glacier. Frontiers in Ecology and Evolution, 9, 584872. https://doi.org/10.3389/fevo.2021.584872

5. Badenes-Pérez, F. R. (2022). Plant-Insect Interactions. Plants, 11(9), 1140. https://doi.org/10.3390/plants11091140

6. Bates, D., Mächler, M., Bolker, B., & Walker, S. (2015). Fitting Linear Mixed-Effects Models Using **lme4**. Journal of Statistical Software, 67(1). https://doi.org/10.18637/jss.v067.i01

7. Bersier, L.-F., Banašek-Richter, C., & Cattin, M.-F. (2002). QUANTITATIVE DESCRIPTORS OF FOOD-WEB MATRICES. Ecology, 83(9), 2394–2407. https://doi.org/10.1890/0012-9658(2002)083[2394:QDOFWM]2.0.CO;2

8. Blüthgen, N., Fründ, J., Vázquez, D. P., & Menzel, F. (2008). WHAT DO INTERACTION NETWORK METRICS TELL US ABOUT SPECIALIZATION AND BIOLOGICAL TRAITS. Ecology, 89(12), 3387–3399. https://doi.org/10.1890/07-2121.1

9. Blüthgen, N., Menzel, F., Blüthgen, N. (2006). Measuring specialization in species interaction networks. BMC Ecology, 6:9. doi:10.1186/1472-6785-6-9

10. Bolker, B. M., Brooks, M. E., Clark, C. J., Geange, S. W., Poulsen, J. R., Stevens, M. H. H., & White, J.-S. S. (2008). Generalized linear mixed models: A practical guide for ecology and evolution. Trends in Ecology & Evolution, 24(3), 127–135. https://doi.org/10.1016/j.tree.2008.10.008

11. Bonelli, M., Eustacchio, E., Avesani, D., Michelsen, V., Falaschi, M., Caccianiga, M., Gobbi, M., & Casartelli, M. (2022). The Early Season Community of Flower-Visiting Arthropods in a High-Altitude Alpine Environment. Insects, 13(4), 393. https://doi.org/10.3390/insects13040393

12. Brighenti, S., Tolotti, M., Bruno, M. C., Wharton, G., Pusch, M. T., & Bertoldi, W. (2019). Ecosystem shifts in Alpine streams under glacier retreat and rock glacier thaw: A review. Science of The Total Environment, 675, 542–559. https://doi.org/10.1016/j.scitotenv.2019.04.221

13. Brochmann, C., Gizaw, A., Chala, D., Kandziora, M., Eilu, G., Popp, M., Pirie, M. D., & Gehrke, B. (2022). History and evolution of the afroalpine flora: In the footsteps of Olov Hedberg. Alpine Botany, 132(1), 65–87. https://doi.org/10.1007/s00035-021-00256-9

14. Brooks, M., Bolker, B., Kristensen, K., Maechler, M., Magnusson, A., McGillycuddy, M., Skaug, H., Nielsen, A., Berg, C., Bentham, K. van, Sadat, N., Lüdecke, D., Lenth, R., O’Brien, J., Geyer, C.J., Jagan, M., Wiernik, B., Stouffer, D.B. (2022). (n.d.). Package *glmmTMB*.

15. Caccianiga, M., Luzzaro, A., Pierce, S., Ceriani, R. M., & Cerabolini, B. (2006). The functional basis of a primary succession resolved by CSR classification. Oikos, 112(1), 10–20. https://doi.org/10.1111/j.0030-1299.2006.14107.x

16. Calatayud, P.-A., Sauvion, N., Thiéry, D., Rebaudo, F., & Jacquin-Joly, E. (2018). Plant-Insect Interactions. In P.-A. Calatayud, N. Sauvion, D. Thiéry, F. Rebaudo, & E. Jacquin-Joly, Ecology. Oxford University Press. https://doi.org/10.1093/obo/9780199830060-0193

17. Cauvy-Fraunié, S., & Dangles, O. (2019). A global synthesis of biodiversity responses to glacier retreat. Nature Ecology & Evolution, 3(12), 1675–1685. https://doi.org/10.1038/s41559-019-1042-8

18. Chapin, F. S., Walker, L. R., Fastie, C. L., & Sharman, L. C. (1994). Mechanisms of Primary Succession Following Deglaciation at Glacier Bay, Alaska. Ecological Monographs, 64(2), 149–175. https://doi.org/10.2307/2937039

19. Chambers, J.M., Hastie, T.J. (1992). *Statistical Models in S*. Wadsworth & Brooks/Cole, Pacific Grove, California. Linear model.

20. Curry, A. M., Cleasby, V., & Zukowskyj, P. (2005). Paraglacial response of steep, sediment-mantled slopes to post-‘Little Ice Age’ glacier recession in the central Swiss Alps. Journal of Quaternary Science, 21(3), 211–225. https://doi.org/10.1002/jqs.954

21. Dormann, C. F., Fruend, J., Gruber, B., Beckett, S., Devoto, M., Felix, G.M.F., Iriondo, J.M., Opsahl, T., Pinheiro, R.B.P., Strauss, R., Diego P. V. (2022). Package *bipartite*.

22. Dormann, C. F., Gruber, B., & Fründ, J. (2008). Introducing the bipartite Package: Analysing Ecological Networks. 8.

23. Dullinger, S., Gattringer, A., Thuiller, W., Moser, D., Zimmermann, N. E., Guisan, A., Willner, W., Plutzar, C., Leitner, M., Mang, T., Caccianiga, M., Dirnböck, T., Ertl, S., Fischer, A., Lenoir, J., Svenning, J.-C., Psomas, A., Schmatz, D. R., Silc, U., … Hülber, K. (2012). Extinction debt of high-mountain plants under twenty-first-century climate change. Nature Climate Change, 2(8), 619–622. https://doi.org/10.1038/nclimate1514

24. Dunne, J. A. (n.d.) (2006). The Network Structure of Food Webs.

25. Dunne, J. A., Williams, R. J., & Martinez, N. D. (2002a). Network structure and biodiversity loss in food webs: Robustness increases with connectance. Ecology Letters, 5(4), 558–567. https://doi.org/10.1046/j.1461-0248.2002.00354.x

26. Dunne, J. A., Williams, R. J., & Martinez, N. D. (2002b). Food-web structure and network theory: The role of connectance and size. Proceedings of the National Academy of Sciences, 99(20), 12917–12922. https://doi.org/10.1073/pnas.192407699

27. Erschbamer, B. (2007). Winners and Losers of Climate Change in a Central Alpine Glacier Foreland. Arctic, Antarctic, and Alpine Research, 39(2), 237–244. https://doi.org/10.1657/1523-0430(2007)39[237:WALOCC]2.0.CO;2

28. Erschbamer, B., & Caccianiga, M. S. (2016). Glacier Forelands: Lessons of Plant Population and Community Development. In F. M. Cánovas, U. Lüttge, & R. Matyssek (Eds.), Progress in Botany Vol. 78 (Vol. 78, pp. 259–284). Springer International Publishing. https://doi.org/10.1007/124_2016_4

29. 29. Fedor, P., & Zvaríková, M. (2019). Biodiversity Indices. In Encyclopedia of Ecology (pp. 337–346). Elsevier. https://doi.org/10.1016/B978-0-12-409548-9.10558-5

30. Fell, S. C., Carrivick, J. L., & Brown, L. E. (2017). The Multitrophic Effects of Climate Change and Glacier Retreat in Mountain Rivers. BioScience, 67(10), 897–911. https://doi.org/10.1093/biosci/bix107

31. Ficetola, G. F., Marta, S., Guerrieri, A., Gobbi, M., Ambrosini, R., Fontaneto, D., Zerboni, A., Poulenard, J., Caccianiga, M., & Thuiller, W. (2021). Dynamics of Ecological Communities Following Current Retreat of Glaciers. *Annual Review of Ecology*, Evolution, and Systematics, 52(1), 405–426. https://doi.org/10.1146/annurev-ecolsys-010521-040017

32. Fijen, T.p.M., Scheper, J.A., Boom, T.M., Janssen, N., Raemakers, I., Kleijn, D. (2018). Insect pollination is at least as important for marketable crop yield as plant quality in a seed crop. Ecology Letters, 21:1704–1713. doi: 10.1111/ele.13150

33. Franzén, M., Dieker, P., Schrader, J., & Helm, A. (2019). Rapid plant colonization of the forelands of a vanishing glacier is strongly associated with species traits. Arctic, Antarctic, and Alpine Research, 51(1), 366–378. https://doi.org/10.1080/15230430.2019.1646574

34. Frenot, Y., Gloaguen, J. C., Cannavacciuolo, M., & Bellido, A. (1998). Primary succession on glacier forelands in the subantarctic Kerguelen Islands. Journal of Vegetation Science, 9(1), 75–84. https://doi.org/10.2307/3237225

35. Gibson, R. H., Knott, B., Eberlein, T., & Memmott, J. (2011). Sampling method influences the structure of plant-pollinator networks. Oikos, 120(6), 822–831. https://doi.org/10.1111/j.1600-0706.2010.18927.x

36. Gilbert, A. J. (2009). Connectance indicates the robustness of food webs when subjected to species loss. Ecological Indicators, 9(1), 72–80. https://doi.org/10.1016/j.ecolind.2008.01.010

37. Giron, D., Dubreuil, G., Bennett, A., Dedeine, F., Dicke, M., Dyer, L. A., Erb, M., Harris, M. O., Huguet, E., Kaloshian, I., Kawakita, A., Lopez-Vaamonde, C., Palmer, T. M., Petanidou, T., Poulsen, M., Sallé, A., Simon, J.-C., Terblanche, J. S., Thiéry, D., … Pincebourde, S. (2018). Promises and challenges in insect-plant interactions. Entomologia Experimentalis et Applicata, 166(5), 319–343. https://doi.org/10.1111/eea.12679

38. Gobbi, M., Ambrosini, R., Casarotto, C., Diolaiuti, G., Ficetola, G. F., Lencioni, V., Seppi, R., Smiraglia, C., Tampucci, D., Valle, B., & Caccianiga, M. (2021a). Vanishing permanent glaciers: Climate change is threatening a European Union habitat (Code 8340) and its poorly known biodiversity. Biodiversity and Conservation, 30(7), 2267–2276. https://doi.org/10.1007/s10531-021-02185-9

39. Gobbi, M., & Lencioni, V. (2021b). Glacial Biodiversity: Lessons from Ground-dwelling and Aquatic Insects. In M. Kanao, D. Godone, & N. Dematteis (Eds.), Glaciers and the Polar Environment. IntechOpen. https://doi.org/10.5772/intechopen.92826

40. Grange, M. C., Munoz, F., Moretti, M., Varona-Y-Varona, S., Renaud, J., Colace, M.-P., Gueguen, M., & Gallien, L. (2021). Designing sampling protocols for plant-pollinator interactions—Timing, meteorology, flowering variations and failed captures matter. Botany Letters, 168(3), 324–332. https://doi.org/10.1080/23818107.2021.1964596

41. Hågvar, S., Gobbi, M., Kaufmann, R., Ingimarsdóttir, M., Caccianiga, M., Valle, B., Pantini, P., Fanciulli, P. P., & Vater, A. (2020). Ecosystem Birth near Melting Glaciers: A Review on the Pioneer Role of Ground-Dwelling Arthropods. Insects, 11(9), 644. https://doi.org/10.3390/insects11090644

42. Hågvar, S. (2012). Primary Succession in Glacier Forelands: How Small Animals Conquer New Land Around Melting Glaciers. In S. Young (Ed.), International Perspectives on Global Environmental Change. InTech. https://doi.org/10.5772/26536

43. Hanusch, M., He, X., Ruiz-Hernández, V., & Junker, R. R. (2022). Succession comprises a sequence of threshold-induced community assembly processes towards multidiversity. Communications Biology, 5(1), 424. https://doi.org/10.1038/s42003-022-03372-2

44. Heleno, R., Devoto, M., & Pocock, M. (2012). Connectance of species interaction networks and conservation value: Is it any good to be well connected? Ecological Indicators, 14(1), 7–10. https://doi.org/10.1016/j.ecolind.2011.06.032

45. Huss, M., Bookhagen, B., Huggel, C., Jacobsen, D., Bradley, R. S., Clague, J. J., Vuille, M., Buytaert, W., Cayan, D. R., Greenwood, G., Mark, B. G., Milner, A. M., Weingartner, R., & Winder, M. (2017). Toward mountains without permanent snow and ice: mountains without permanent snow and ice. Earth’s Future, 5(5), 418–435. https://doi.org/10.1002/2016EF000514

46. Inouye, D. W. (2020). Effects of climate change on alpine plants and their pollinators. Annals of the New York Academy of Sciences, 1469(1), 26–37. https://doi.org/10.1111/nyas.14104

47. Junker, R. R., Hanusch, M., He, X., Ruiz-Hernández, V., Otto, J.-C., Kraushaar, S., Bauch, K., Griessenberger, F., Ohler, L.-M., & Trutschnig, W. (2020). Ödenwinkel: An Alpine platform for observational and experimental research on the emergence of multidiversity and ecosystem complexity. Web Ecology, 20(2), 95–106. https://doi.org/10.5194/we-20-95-2020

48. Kaufmann R, Raffl C. 2002. Diversity in primary succession: the chronosequence of a glacier foreland. In Mountain Diversity: A Global Assessment, ed. C Körner, EM Spehn, pp. 177–90. Boca Raton, FL: CRC Press

49. Kiester, A. R. (2013). Species Diversity, Overview. In Encyclopedia of Biodiversity (pp. 706–714). Elsevier. https://doi.org/10.1016/B978-0-12-384719-5.00133-7

50. Konopiński, M. K. (2020). Shannon diversity index: A call to replace the original Shannon’s formula with unbiased estimator in the population genetics studies. PeerJ, 8, e9391. https://doi.org/10.7717/peerj.9391

51. Lambiel, C., Maillard, B., Kummert, M., & Reynard, E. (2016). Geomorphology of the Hérens valley (Swiss Alps). Journal of Maps, 12(1), 160–172. https://doi.org/10.1080/17445647.2014.999135

52. Lefebvre, V., Villemant, C., Fontaine, C., & Daugeron, C. (2018). Altitudinal, temporal and trophic partitioning of flower-visitors in Alpine communities. Scientific Reports, 8(1), 4706. https://doi.org/10.1038/s41598-018-23210-y

53. Llambí, L. D., Melfo, A., Gámez, L. E., Pelayo, R. C., Cárdenas, M., Rojas, C., Torres, J. E., Ramírez, N., Huber, B., & Hernández, J. (2021). Vegetation Assembly, Adaptive Strategies and Positive Interactions During Primary Succession in the Forefield of the Last Venezuelan Glacier. Frontiers in Ecology and Evolution, 9, 657755. https://doi.org/10.3389/fevo.2021.657755

54. Losapio, G., Cerabolini, B. E. L., Maffioletti, C., Tampucci, D., Gobbi, M., & Caccianiga, M. (2021a). The Consequences of Glacier Retreat Are Uneven Between Plant Species. Frontiers in Ecology and Evolution, 8, 616562. https://doi.org/10.3389/fevo.2020.616562

55. Losapio, G., Norton Hasday, E., Espadaler, X., Germann, C., Ortiz-Sánchez, F. J., Pont, A., Sommaggio, D., & Schöb, C. (2021b). Facilitation and biodiversity jointly drive mutualistic networks. Journal of Ecology, 109(5), 2029–2037. https://doi.org/10.1111/1365-2745.13593

56. Losapio, G., Schmid, B., Bascompte, J., Michalet, R., Cerretti, P., Germann, C., Haenni, J., Neumeyer, R., Ortiz-Sánchez, R.J., C. Pont, A., Rousse, P., Schmid, J., Sommaggio, D., Schöb, C. (2020). An experimental approach to assessing the impact of ecosystem engineers on biodiversity and ecosystem functions. Ecology, 102(2), e03243. https://doi.org/10.1002/ecy.3243

57. Losapio, G., Fortuna, M. A., Bascompte, J., Schmid, B., Michalet, R., Neumeyer, R., Castro, L., Cerretti, P., Germann, C., Haenni, J., Klopfstein, S., Ortiz-Sanchez, F. J., Pont, A. C., Rousse, P., Schmid, J., Sommaggio, D., & Schöb, C. (2019). Plant interactions shape pollination networks via nonadditive effects. Ecology, 100(3). https://doi.org/10.1002/ecy.2619

58. Losapio, G., de la Cruz, M., Escudero, A., Schmid, B., & Schöb, C. (2018). The assembly of a plant network in alpine vegetation. Journal of Vegetation Science, 29(6), 999– 1006. https://doi.org/10.1111/jvs.12681

59. Losapio, G., Gobbi, M., Marano, G., Avesani, D., Boracchi, P., Compostella, C., Pavesi, M., Schöb, C., Seppi, R., Sommaggio, D., Zanetti, A., & Caccianiga, M. (2016). Feedback effects between plant and flower-visiting insect communities along a primary succession gradient. Arthropod-Plant Interactions, 10(6), 485–495. https://doi.org/10.1007/s11829-016-9444-x

60. Losapio, G., Jordánc, F., Caccianiga, M., Gobbi, M. (2015). Structure-dynamic relationship of plant–insect networks along a primary succession gradient on a glacier foreland. Ecological Modelling 314: 73–79. http://dx.doi.org/10.1016/j.ecolmodel.2015.07.014

61. Marta, S., Azzoni, R. S., Fugazza, D., Tielidze, L., Chand, P., Sieron, K., Almond, P., Ambrosini, R., Anthelme, F., Alviz Gazitúa, P., Bhambri, R., Bonin, A., Caccianiga, M., Cauvy-Fraunié, S., Ceballos Lievano, J. L., Clague, J., Cochachín Rapre, J. A., Dangles, O., Deline, P., … Ficetola, G. F. (2021). The Retreat of Mountain Glaciers since the Little Ice Age: A Spatially Explicit Database. Data, 6(10), 107. https://doi.org/10.3390/data6100107

62. Marzeion, B., Cogley, J. G., Richter, K., & Parkes, D. (2014). Attribution of global glacier mass loss to anthropogenic and natural causes. Science, 345(6199), 919–921. https://doi.org/10.1126/science.1254702

63. Martínez-Núñez, C., Kleijn, D., Ganuza, C., Heupink, D., Raemakers, I., Vertommen, W., & Fijen, T. P. M. (2022). Temporal and spatial heterogeneity of semi-natural habitat, but not crop diversity, is correlated with landscape pollinator richness. Journal of Applied Ecology, 59(5), 1258–1267. https://doi.org/10.1111/1365-2664.14137

64. Matthews, J.A., (1992). The ecology of recently-deglaciated terrain. A geoecological approach to glacier forelands and primary succession. Antarctic Science 4(4): 484–485. https://doi.org/10.1017/S0954102092260715

65. McCabe, L. M., & Cobb, N. S. (2021). From Bees to Flies: Global Shift in Pollinator Communities Along Elevation Gradients. Frontiers in Ecology and Evolution, 8, 626124. https://doi.org/10.3389/fevo.2020.626124

66. Nicolussi, K., Le Roy, M., Schlüchter, C., Stoffel, M., & Wacker, L. (2022). The glacier advance at the onset of the Little Ice Age in the Alps: New evidence from Mont Miné and Morteratsch glaciers. The Holocene, 32(7), 624–638. https://doi.org/10.1177/09596836221088247

67. Oksanen J, Blanchet FG, Friendly M, Kindt R, Legendre P, McGlinn D, Minchin PR, O’Hara RB, Simpson GL, Solymos P, Stevens MHH, Szoecs E, Wagner H. 2019. vegan: community ecology package. R package version 2.5-6. Available at https://cran.r-project.org/package=vegan.

68. Paul, F., Rastner, P., Azzoni, R. S., Diolaiuti, G., Fugazza, D., Le Bris, R., Nemec, J., Rabatel, A., Ramusovic, M., Schwaizer, G., & Smiraglia, C. (2020). Glacier shrinkage in the Alps continues unabated as revealed by a new glacier inventory from Sentinel-2. Earth System Science Data, 12(3), 1805–1821. https://doi.org/10.5194/essd-12-1805-2020

69. Pimm, S. L. (1984a). The complexity and stability of ecosystems. Nature, 307(5949), 321–326. https://doi.org/10.1038/307321a0

70. Portet, S. (2020). A primer on model selection using the Akaike Information Criterion. Infectious Disease Modelling, 5, 111–128. https://doi.org/10.1016/j.idm.2019.12.010

71. Robinson, S. V. J., Losapio, G., & Henry, G. H. R. (2018). Flower-power: Flower diversity is a stronger predictor of network structure than insect diversity in an Arctic plant– pollinator network. Ecological Complexity, 36, 1–6. https://doi.org/10.1016/j.ecocom.2018.04.005

72. Roe, G. H., Baker, M. B., & Herla, F. (2017). Centennial glacier retreat as categorical evidence of regional climate change. Nature Geoscience, 10(2), 95–99. https://doi.org/10.1038/ngeo2863

73. Rosero, P., Crespo-Pérez, V., Espinosa, R., Andino, P., Barragán, Á., Moret, P., Gobbi, M., Ficetola, G. F., Jaramillo, R., Muriel, P., Anthelme, F., Jacobsen, D., Dangles, O., Condom, T., Gielly, L., Poulenard, J., Rabatel, A., Basantes, R., Cáceres Correa, B., & Cauvy-Fraunié, S. (2021). Multi-taxa colonisation along the foreland of a vanishing equatorial glacier. Ecography, 44(7), 1010–1021. https://doi.org/10.1111/ecog.05478

74. Sommer, C., Malz, P., Seehaus, T. C., Lippl, S., Zemp, M., & Braun, M. H. (2020). Rapid glacier retreat and downwasting throughout the European Alps in the early 21st century. Nature Communications, 11(1), 3209. https://doi.org/10.1038/s41467-020-16818-0

75. Stibal, M., Bradley, J. A., Edwards, A., Hotaling, S., Zawierucha, K., Rosvold, J., Lutz, S., Cameron, K. A., Mikucki, J. A., Kohler, T. J., Šabacká, M., & Anesio, A. M. (2020). Glacial ecosystems are essential to understanding biodiversity responses to glacier retreat. Nature Ecology & Evolution, 4(5), 686–687. https://doi.org/10.1038/s41559-020-1163-0

76. Tampucci, D., Gobbi, M., Boracchi, P., Cabrini, E., Compostella, C., Mangili, F., Marano, G., Pantini, P., & Caccianiga, M. (2015). Plant and arthropod colonisation of a glacier foreland in a peripheral mountain range. Biodiversity, 16(4), 213–223. https://doi.org/10.1080/14888386.2015.1117990

77. Trunschke, J., & Stöcklin, J. (2017). Plasticity of flower longevity in alpine plants is increased in populations from high elevation compared to low elevation populations. Alpine Botany, 127(1), 41–51. https://doi.org/10.1007/s00035-016-0176-4

78. Valle, B., di Musciano, M., Gobbi, M., Bonelli, M., Colonnelli, E., Gardini, G., Migliorini, M., Pantini, P., Zanetti, A., Berrilli, E., Frattaroli, A. R., Fugazza, D., Invernizzi, A., & Caccianiga, M. (2022a). Biodiversity and ecology of plants and arthropods on the last preserved glacier of the Apennines Mountain chain (Italy). The Holocene, 32(8), 853–865. https://doi.org/10.1177/09596836221096292

79. Valle, B., Gobbi, M., Tognetti, M., Borgatti, M. S., Compostella, C., Pantini, P., & Caccianiga, M. (2022b). Glacial biodiversity of the southernmost glaciers of the European Alps (Clapier and Peirabroc, Italy). Journal of Mountain Science, 19(8), 2139–2159. https://doi.org/10.1007/s11629-022-7331-8

80. Vitasse, Y., Ursenbacher, S., Klein, G., Bohnenstengel, T., Chittaro, Y., Delestrade, A., Monnerat, C., Rebetez, M., Rixen, C., Strebel, N., Schmidt, B. R., Wipf, S., Wohlgemuth, T., Yoccoz, N. G., & Lenoir, J. (2021). Phenological and elevational shifts of plants, animals and fungi under climate change in the E uropean A lps. Biological Reviews, 96(5), 1816–1835. https://doi.org/10.1111/brv.12727

81. Whittaker, R. J. (1993). Plant population patterns in a glacier foreland succession: Pioneer herbs and later-colonizing shrubs. Ecography, 16(2), 117–136. https://doi.org/10.1111/j.1600-0587.1993.tb00064.x

82. Zemp, M., Frey, H., Gärtner-Roer, I., Nussbaumer, S. U., Hoelzle, M., Paul, F., Haeberli, W., Denzinger, F., Ahlstrøm, A. P., Anderson, B., Bajracharya, S., Baroni, C., Braun, L. N., Cáceres, B. E., Casassa, G., Cobos, G., Dávila, L. R., Delgado Granados, H., Demuth, M. N., … Vincent, C. (2015). Historically unprecedented global glacier decline in the early 21st century. Journal of Glaciology, 61(228), 745–762. https://doi.org/10.3189/2015JoG15J017

83. Zemp, M., Huss, M., Thibert, E., Eckert, N., McNabb, R., Huber, J., Barandun, M., Machguth, H., Nussbaumer, S. U., Gärtner-Roer, I., Thomson, L., Paul, F., Maussion, F., Kutuzov, S., & Cogley, J. G. (2019). Global glacier mass changes and their contributions to sea-level rise from 1961 to 2016. Nature, 568(7752), 382–386. https://doi.org/10.1038/s41586-019-1071-0

84. https://www.infoflora.ch

85. https://www.inaturalist.org/projects/pollinator-diversity-at-ferpecle-glacier-ecosystems

86. https://map.geo.admin.ch

